# The oscillation of mitotic kinase governs cell cycle latches in mammalian cells

**DOI:** 10.1101/2023.05.19.541462

**Authors:** Calin-Mihai Dragoi, Alexis R. Barr, John J. Tyson, Béla Novák

## Abstract

The mammalian cell cycle alternates between two phases: S-G2-M with high levels of A- and B-type cyclin-dependent kinases (CycA,B:CDK); and G1 with persistent degradation of CycA,B by Cdh1-activated APC/C (anaphase promoting complex/cyclosome). Because CDKs phosphorylate and inactivate Cdh1, these two phases are mutually exclusive. This ‘toggle switch’ is flipped from G1 to S by cyclin-E (CycE:CDK), which is not degraded by Cdh1:APC/C; and from M to G1 by Cdc20:APC/C, which is not inactivated by CycA,B:CDK. After flipping the switch, cyclin E is degraded and Cdc20:APC/C is inactivated. Combining mathematical modelling with single-cell timelapse imaging, we show that dysregulation of CycB:CDK disrupts strict alternation of the G1-S and M-G1 switches. Inhibition of CycB:CDK results in Cdc20-independent Cdh1 ‘endocycles’, and sustained activity of CycB:CDK drives Cdh1-independent Cdc20 endocycles. Our model provides one mechanistic explanation for how whole genome doubling can arise, a common event in tumorigenesis that can drive tumour evolution.

## INTRODUCTION

The eukaryotic cell cycle is the repetitive process of DNA synthesis (chromosome replication, S), mitosis (alignment of the replicated chromosomes on the mitotic spindle, M), anaphase (separation of the sister chromatids to opposite poles of the spindle, A), telophase (formation of daughter nuclei, each containing a full complement of unreplicated chromosomes, T), and cell division (separation into two, genetically identical daughter cells, CD, Fig.1A). This cycle of DNA replication and chromosome partitioning runs in parallel to cell growth, whereby all other essential components of the cell (proteins, lipids, polysaccharides, organelles) are amplified and divided more-or-less evenly between the newborn daughter cells. The growth and division processes are balanced, in the long run, so that a proliferating cell population maintains stable size and DNA distributions ^1^.

**Figure 1.**
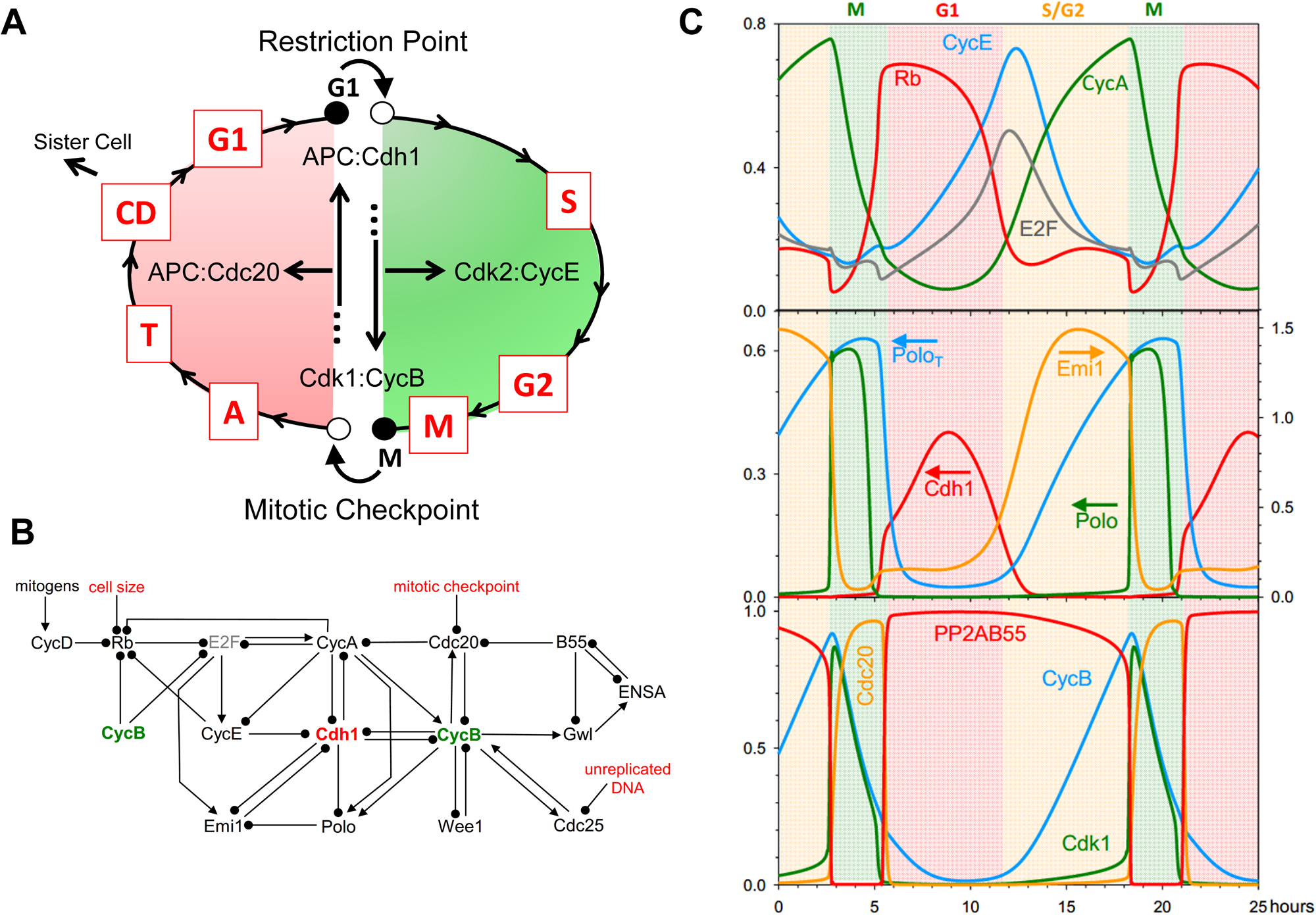
A model of mammalian cell cycle controls. **(A)** Conceptual framework. A newborn cell arrests in G1 phase (unreplicated chromosomes) at a stable steady state (●), which we call **G1**. Growth factors, integrated at the Restriction Point (RP), destabilize **G1** (o) and induce the cell to enter S/G2/M phase (green), replicating its chromosomes and eventually arresting in mitosis at a different stable steady state called **M**, while the replicated chromosomes are coming into alignment on the mitotic spindle. When the spindle is properly assembled and all chromosomes are properly aligned, the mitotic checkpoint is lifted, destabilizing **M** (●→o) and allowing the cell to exit mitosis (red phase: M→A→T), divide (CD) and return to the **G1** stable state. These events are coordinated by a complex protein interaction network, whose principal components are displayed inside the cycle. **(B)** An influence diagram summarizing mammalian cell cycle controls. Arrow-heads indicate ‘activation’ or ‘synthesis’; black-dots indicate ‘inactivation’ or ‘degradation’. Cdh1 and CycB play central roles in the control system. At the **G1** steady state, Cdh1 and Rb are active, E2F is inactive, the cyclins (A, B, E and D) are low, as are Emi1 and Polo. At the **M** steady state, Cdh1 is inactive, and CycA, CycB and Polo are active. This diagram is converted into a set of nonlinear ordinary differential equations in the Materials and Methods. **(C)** Limit-cycle oscillations of the model when all checkpoints are removed. The model ODEs are simulated numerically for the parameter values given in Table 1, and selected variables are plotted as functions of time (in hours). The phases of the cell cycle are color coded: G1 (pink), S/G2 (yellow) and M (green). Notice that Rb and Cdh1 activities are high in G1 phase; CycE and E2F activities peak at the G1/S transition; Emi1 and CycA are high in G2 phase; ‘Cdk1 activity’ (i.e., active Cdk1:CycB) and Polo peak as the simulated cell enters mitosis, and Cdc20 peaks as the cell exits mitosis and returns to G1 phase. Meanwhile, PP2A:B55 activity is high throughout G1/S/G2 and low only when Cdk1 activity is high. Because no checkpoint controls are operational in this simulation, the cell cycle time-courses do not pause at the stable steady states (**G1** and **M**) in panel A. In the middle panel, arrows indicate the corresponding y-axis for dynamic variables.

**Table 1:**
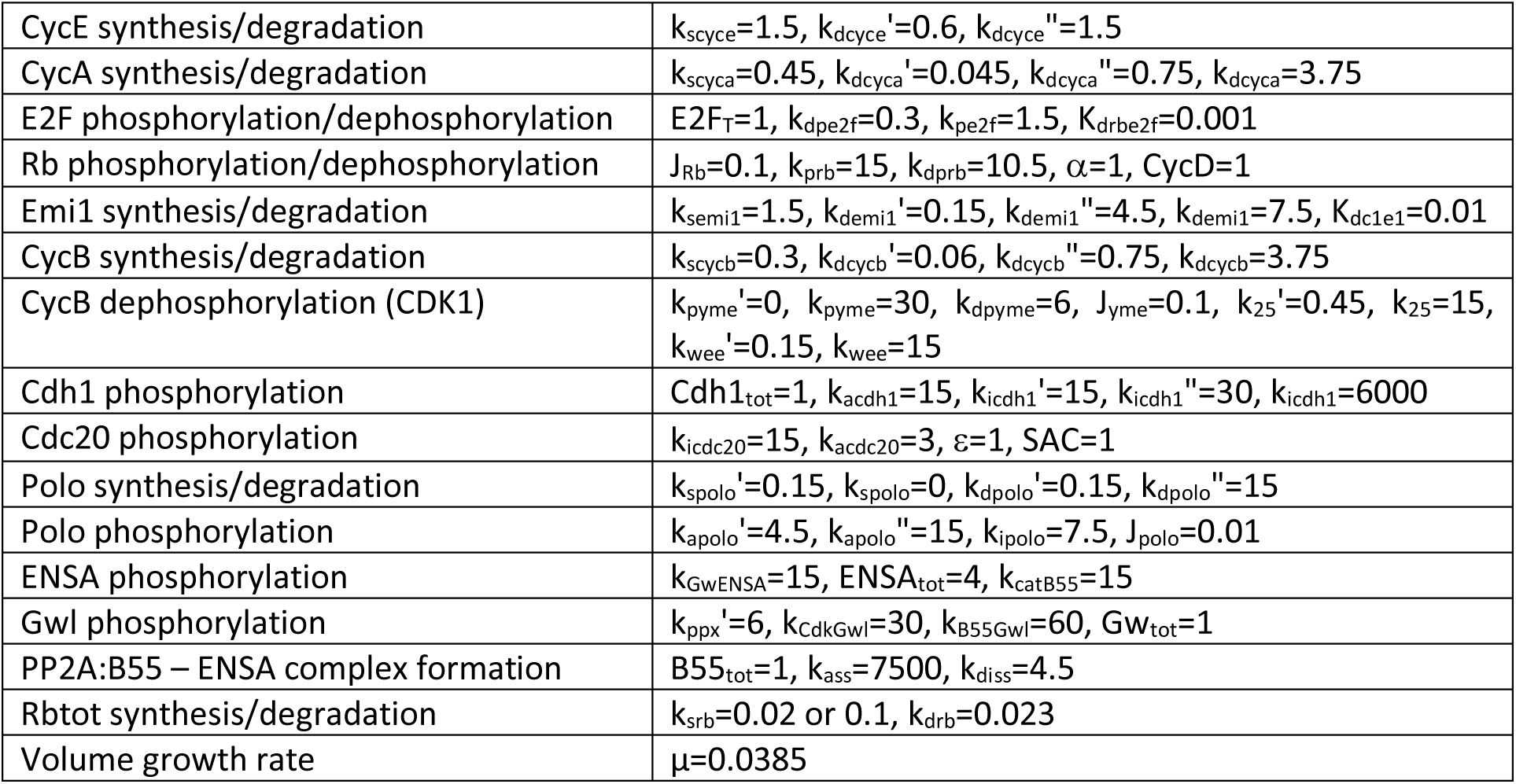
Kinetic parameters of the mammalian cell model. Rate constants (k’s) have a dimension of h-1 while other parameters are dimensionless.

Eukaryotic cells coordinate growth and division at cell cycle ‘checkpoints.’ Animal cells normally arrest soon after birth (in G1 phase of the cycle, with unreplicated chromosomes) in a stable ‘pause’ state, known as quiescence. To re-enter a new round of growth and division, cells must first pass the Restriction Point (RP). To pass the RP and progress into S-phase (DNA replication), a cell must receive ‘permission’ in the form of extracellular growth factors, that disengage the ‘brakes’ holding the cell prior to the RP ^2^. After passing the RP, the cell replicates its DNA and enters mitosis. A second crucial checkpoint, the spindle assembly checkpoint, arrests the cell in mitosis until all the replicated chromosomes are properly aligned on the mitotic spindle ^3^. Then, and only then, the cell receives permission to initiate anaphase and partition the sister chromatids to daughter nuclei.

These checkpoints and transitions are implemented by an exceedingly complex network of interacting genes and proteins ^4^. In earlier publications ^5–7^ we have studied this network in detail for budding yeast cells (*Saccharomyces cerevisiae*). We proposed that—at its core—the cell cycle is an alternation between two fundamental phases: G1 (not committed to DNA replication and cell division) and S/G2/M (in progress toward mitosis and cell division). Each phase is attracted to a <u>stable steady state</u> of the underlying molecular regulatory network; let us denote them **G1** and **M**. S/G2/M phase is characterized by rising activity of B-type cyclin-dependent kinases (CDKs), driving S phase and entry into mitosis. S/G2/M phase ultimately terminates at **M**, a stable steady state characterized by high CDK activity and negligible activities of the CDK antagonists that are prevalent in G1. The uncommitted phase is characterized by a stable steady state, **G1**, of low CDK activity and high antagonist activities.

In budding yeast, the principal CDK-antagonists are the Anaphase Promoting Complex/Cyclosome (APC/C), which promotes polyubiquitination of B-type cyclins (Clb1-5) and their subsequent degradation by proteasomes, and cyclin-dependent kinase inhibitors (CKIs), which bind to and inhibit the B-type CDKs ^8^. In G1 phase, APC/C activity is directed towards Clb1-5 by a targeting subunit called Cdh1. Hence, we characterize **G1** as a steady state with high activities of both APC/C:Cdh1 and CKIs. The B-type CDKs and their antagonists are mutually inhibitory: not only do the antagonists suppress CDK activity, but the CDKs phosphorylate both Cdh1 (Cdh1-P is inactive) and CKI (CKI-P is rapidly degraded by the Skp/Cullin/F-box (SCF) polyubiquitination pathway) ^8^. The mutual inhibition between B-type CDKs and their antagonists is the source of the coexisting, stable steady states (**M** and **G1)** of the underlying cell-cycle control system in yeast ^5–7^.

Although we originally presented this concept of cell cycle regulation for budding yeast, it is applicable to eukaryotic organisms in general, because B-type CDKs are a universal feature of entry into mitosis ^9^, and their opposition by APC/C:Cdh1 and stoichiometric CKIs is a universal feature of G1-arrest in eukaryotic cells. In this paper, we focus on the control system in mammalian cells. As suggested by Fig. 1A, the coexisting stable steady states (**G1** and **M**) of the underlying bistable switch force the cell to follow a distinctive loop of cell-cycle events governed by two characteristic transitions: from G1 into S/G2/M as the Restriction Point (RP) is lifted, and from M into A/T/CD/G1 as the Mitotic Checkpoint (MC) is lifted (Fig. 1A). At these transitions, the cell executes one of the two crucial events of the chromosome cycle: passing from G1 into S/G2/M, chromosomes are replicated and brought into alignment on the mitotic spindle; and passing from M into A/T/CD/G1, the sister chromatids are partitioned to two daughter cells.

These two transitions are fundamentally irreversible because of a ‘latching’ property of the bistable switch. At RP, the **G1** steady state becomes unstable (●→o), and the cell enters S/G2/M by upregulating Cdk2:CycE, which promotes the rise of Cdk2:CycA (involved in DNA replication) and Cdk1:CycB (mitotic CDK activity). As CycA- and CycB-dependent kinases rise, CycE is phosphorylated and degraded by the SCF pathway (negative feedback ^10^). As CycE-dependent kinase activity drops, the control system is captured by the stable steady state **M** (● in Fig. 1A). At RP, the ‘G1 gate’ is opened and CycE pushes the cell into S/G2/M. The negative feedback loop acts as a ‘spring’ to pull the gate closed, and it ‘latches’ at the stable **M** state. For the cell to divide and return to G1 phase, the MC must destabilize **M** (●→o), causing APC/C:Cdc20 to polyubiquitinate/degrade both Securin and CycB ^11^, which allows sister chromatids to separate and the cell to proceed into A and T. As CycB-dependent kinase activity drops, the APC/C dissociates from Cdc20 and binds to Cdh1 ^12^. The falling activity of APC/C:Cdc20 is the ‘spring’ that pulls the mitotic-exit gate closed and latched at the stable **G1** state.

The irreversible ‘latching’ property of these gates guarantees that a proliferating cell alternates between S phase (DNA replication) and mitosis (accurate partitioning of replicated chromosomes to the two incipient daughter cells). A cell that leaves G1 and enters S phase is captured by the **M** latch. The cell can only divide and return to G1 by destabilizing **M** and getting captured by the **G1** latch.

The alternation between **G1** and **M** is facilitated by ‘helper’ molecules (a starter kinase like CycE:Cdk2 and an exit protein like Cdc20). The helper molecules are regulated by negative feedback mechanisms that inactivate them after the transition is triggered ^6, 7^. The latching behavior requires that the control system alternate between two different steady states: **G1** (low CDK activity and active CDK-antagonists) and **M** (high CDK activity and inactive CDK-antagonists).

In this work, we show that this informal, verbal description of cell-cycle progression is a precise mathematical consequence of the molecular interactions among the CDKs, antagonists and helpers of the mammalian cell cycle control system. Our mathematical model makes interesting predictions about the appearance of ‘endocycles’ (e.g., periodic DNA replication without mitosis, or periodic oscillations of CycB-dependent kinase activity without DNA replication) when the latching gates at **G1** and **M** are compromised.

## RESULTS

### Proposed Mechanism and Mathematical Model

At the heart of our model is APC/C:Cdh1, which is regulated by Cdh1-inhibitory phosphorylations by CycE-, CycA- and CycB-associated CDK activities (Fig. 1B, ^11^). The double-negative interactions between Cdh1 and CycA- and CycB-dependent kinases are fundamental to the alternative stable steady states, **G1** and **M**, of our model. Cdh1 is also regulated by Emi1 (Early Mitotic Inhibitor 1), which is an inhibitory substrate of APC/C:Cdh1 and accounts for a third double-negative feedback loop that renders APC/C:Cdh1 activity bistable ^13^. To leave G1 phase and enter S/G2/M, APC/C:Cdh1 activity must be suppressed, and this is initiated by the transcription factor, E2F. To prevent premature entry into S phase, E2F is inhibited in G1 phase by the retinoblastoma protein (Rb), the primary agent arresting G1 cells at the restriction point. To pass RP, the cell must inactivate Rb by phosphorylation, started by CycD:Cdk4/Cdk6 (CDK activities under the control of a variety of ‘mitogens,’ ‘growth factors’ and ‘anti-growth factors’). Rb phosphorylation releases E2F to induce the synthesis of CycE, CycA and Emi1. CycE and Emi1 combine to drive down Cdh1-dependent degradation of CycA and CycB. Both CycE and CycA can drive the cell into S phase, and phosphorylation of CycE targets it for ubiquitination and degradation by the SCF pathway. CycA also activates the MuvB transcription factor for CycB expression ^14^. Rising activities of CycA and CycB propel the cell through G2 into M phase. Cyclins E, A and B also amplify and prolong the phosphorylation of Rb started by CycD. During S phase, after the burst in E2F-mediated transcription, E2F is inactivated by phosphorylation by CycA- and CycB CDKs ^15^. Another kinase involved in APC/C:Cdh1 regulation is Polo. When Cdh1 activity is low (in S/G2/M), the accumulating Polo-kinase is indirectly activated by CycA and CycB ^16^. Polo is responsible for phosphorylating Emi1, thereby promoting Emi1 degradation before mitotic entry ^17^ and leaving A-type and B-type CDKs the only remaining activities to maintain Cdh1 inactive until mitotic exit.

The other central component of our model is CycB:Cdk1, whose activity drives the cell into mitosis and whose degradation allows the cell to exit mitosis ^9^. CycB:Cdk1 complexes undergo inhibitory tyrosine-phosphorylation on the Cdk1 subunit by Wee1-kinase and dephosphorylation by Cdc25-phosphatases. The abrupt rise of Cdk1 activity at the onset of mitosis is triggered by the positive feedback loops between Cdk1|Wee1 (−|−) and Cdk1|Cdc25 (+|+). (The bistability of this activation process ^18–20^ creates the opportunity for an ‘unreplicated DNA’ checkpoint at the G2/M transition; an important check on genome integrity that we shall not pursue further in this paper). Rising CycB:Cdk1 activity phosphorylates both APC/C and Greatwall-kinase (Gwl). Phosphorylated APC/C rapidly binds to Cdc20, and the active complex (P-APC/C:Cdc20) promotes the degradation of both CycA and CycB. Meanwhile, activated Gwl phosphorylates and activates endosulfine (ENSA), which inhibits the major phosphatase (PP2A:B55) during mitosis ^21, 22^. As long as PP2A:B55 is inhibited, APC/C:Cdc20 actively clears CycA and CycB from the cell; but, as CycB:Cdk1 activity drops, the balance between APC/C phosphorylation and dephosphorylation shifts to favor its dissociation from Cdc20 and association with dephosphorylated Cdh1. These molecular changes drive the cell back to G1 (active APC/C:Cdh1). If Rb is active in the newborn cell, it will arrest at **G1** (the RP).

In the Methods section, we provide a set of ordinary differential equations (ODEs) that describe our proposed mechanism (Fig. 1B) for control of the mammalian cell cycle.

### A Cell-Cycle Clock

We start our analysis of the mathematical model by numerical integration of the ODEs in the absence of checkpoint regulation at RP or MC. With appropriate choice of kinetic parameters, numerical simulations exhibit persistent limit-cycle oscillations, corresponding to an autonomous cell-cycle ‘clock’ (Fig. 1C). As expected, in G1 phase, Cdh1 is active and unphosphorylated Rb is high. As E2F activity rises, CycE is the first E2F target to appear, because it is not degraded by Cdh1. CycE phosphorylates Cdh1 and Rb, causing their activities to drop, allowing CycA and Emi1 to rise, which are hallmarks of the G1/S transition ^13, 23^. The rise of CycB is delayed until CycA activates the MuvB transcription factor complex. As CycB level rises, CycB:Cdk1 is activated by the positive feedback-aided dephosphorylation of Cdk1. High CycB-dependent kinase activity activates Polo and APC/C:Cdc20 and inactivates PP2A:B55 via the Gwl-ENSA pathway. Polo activation causes degradation of Emi1 (the Cdh1 inhibitor), but Cdh1-dependent APC/C activity remains low because high CDK activity phosphorylates Cdh1 and inhibits its association with APC/C. CycB-activated

APC/C:Cdc20 maintains its activity until CycB is almost completely degraded, because the APC/C-inactivating phosphatase (PP2A:B55) is inhibited by ENSA.

### Mapping the Cell Cycle Clock with Bifurcation Curves

The previous section illustrates that without any checkpoint control our model of the mammalian cell cycle exhibits a limit cycle oscillation. To provide insight into this clock mechanism, we turn to bifurcation diagrams. A bifurcation diagram plots the steady state value of a cell cycle regulator as a function of increasing values of a bifurcation parameter. We choose CycE and Cdc20 as bifurcation parameters, because they act as helper molecules for the G1/S and the M/G1 transitions, respectively. To characterize the state of the cell cycle control system, we choose either Cdh1 activity or the level of CycB (mitotic cyclin). Since the changes of the two helper molecules are almost out-of-phase during the cycle (see Fig. 1C), we set Cdc20 = 0 when calculating the bifurcation diagram for CycE and CycE = 0 for the Cdc20 diagram. To be more precise, to calculate the bifurcation diagram with CycE as the parameter, we eliminate the differential equations for both d[CycE]/dt and d[Cdc20]/dt, then set [Cdc20] = 0 and [CycE] = constant everywhere in the remaining ODEs. We then solve for the steady state of the remaining nonlinear ODEs as a function of the value of [CycE], using the bifurcation software AUTO as implemented in XPP ^24^.

The Cdh1 bifurcation diagrams (Fig. 2) show a Z-shaped dependence on each of the helper molecules, CycE and Cdc20. Notice that the G1/S and M/G1 switches are <u>irreversible</u>, in the following sense. To leave G1 (Cdh1 active) and enter S/G2/M (Cdh1 activity falling to ∼0), CycE activity must increase above ∼0.47 to get beyond the ‘nose’ of stable steady states of high Cdh1 activity (see Fig. 2A; the ‘nose’ is called a ‘saddle-node’ bifurcation point). Thereafter, the trajectory drops to the branch of lower steady states, and, as CycE is degraded (as a consequence of CycE- and CycA-dependent phosphorylation and SCF-dependent ubiquitination), the trajectory stops at **M** because it can go no further. To switch back to G1 phase spontaneously, [CycE] would have fall to negative values. Similarly, the M/G1 switch is irreversible because [Cdc20] would have to fall to negative values. For these reasons, spontaneous ‘endocycles’ (G1/S/G1/S/… and M/G1/M/G1/…) are impossible (under the present conditions), and progression through the cell cycle is an irreversible alternation between G1/S and M/G1 transitions, as suggested by the cell cycle trajectory (dotted black curves in Fig. 2).

**Figure 2.**
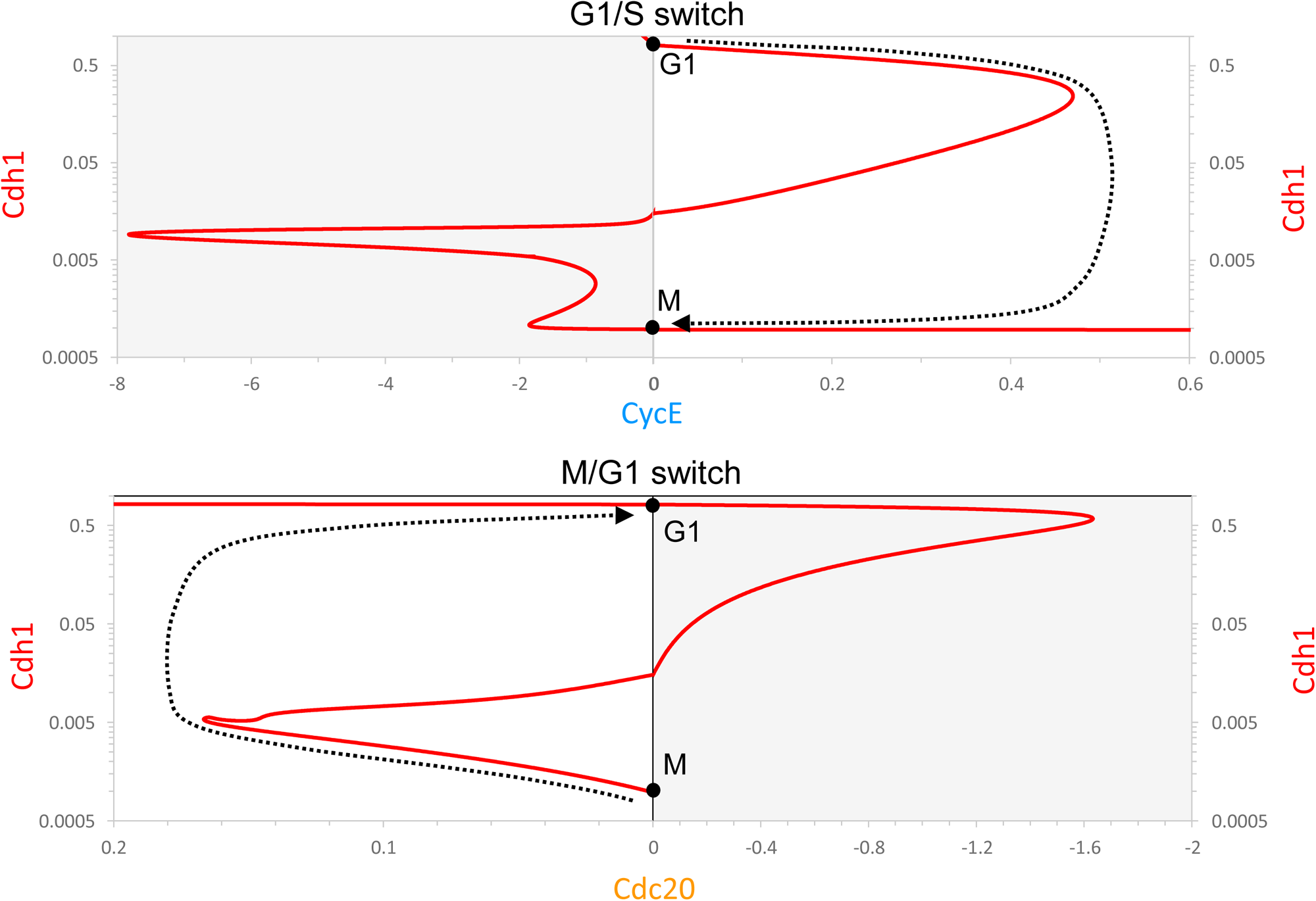
Bifurcation diagrams for Cdh1 activity as a function of CycE or Cdc20. Red curve: steady state activity of APC/C:Cdh1 as a function of [CycE] (top) or [Cdc20] (bottom); dotted black lines: proposed cell-cycle ‘trajectory’ projected onto the bifurcation diagram, based on the negative feedback loops controlling CycE and Cdc20. To calculate the CycE diagram, we set [Cdc20] = 0; for the Cdc20 diagram, we set [CycE] = 0. The **G1** and **M** steady states are marked by ●. Notice that, for the CycE diagram, positive values of [CycE] are plotted to the right and negative values (shaded, which cannot be visited by the system) to the left. For the Cdc20 diagram, the positive and negative values are reversed, for a reason that will soon be apparent.

The corresponding CycB bifurcation diagrams (Fig. S1) are S-shaped, mirroring the Cdh1 curve (Fig. 2), because Cdh1 activity and CycB levels mirror each other. When [CycE] exceeds ∼0.47 (Suppl. Fig. S1A), Cdh1 becomes inactivated and CycB level increases. Since CycE is regulated by a negative feedback loop, its level decreases after the G1/S transition, as CycB is accumulating. As CycE level falls, Cdh1 does not become reactivated, because the reactivation threshold is at a negative value of [CycE]. Both CycA (not shown) and CycB reach stable steady-state values (**M**) as [CycE] → 0.

To reactivate Cdh1, the other helper molecule, APC/C:Cdc20, must be activated above a threshold value of ∼0.17 (Fig. 2B), which leads to the degradation of both CycA and CycB (Suppl. Fig. S1B).

Because APC/C:Cdc20 activity depends upon APC/C phosphorylation by Cdk1:CycB, Cdc20 activity falls as CycB activity falls (with a slight time delay). Cdh1, on the other hand, stays active and keeps CycB at a low steady-state level (**G1**) after Cdc20 inactivation. CycB does not spontaneously reaccumulate, because the CycB reactivation threshold is at negative Cdc20 value (−1.6). In this way, active Cdh1 latches the gate after the cell exits mitosis.

In summary, both G1/S and M/G1 transitions are governed by irreversible bistable switches (‘latching gates’).

To put together a picture of the whole cell cycle, we combine the two half-bifurcation diagrams calculated with CycE and Cdc20 as helper molecules (Fig. 3). Keep in mind that these diagrams are approximations based on our reasonable simplifying assumption that the two helper molecules do not coexist, i.e., Cdc20 and CycE are absent (equal to zero) on the right and left sides, respectively. The combined Cdh1 bifurcation diagram maintains the characteristic Z-shape of the Cdh1 vs CycE and Cdh1 vs Cdc20 diagrams (Fig. 2A). Similarly, the combined CycB bifurcation diagram (Fig. 2B) maintains the S-shape of the diagrams in Fig. S1. According to our model, opening the G1/S gate triggers the transition from **G1** to the alternative **M** steady state and also latches the M/G1 gate by inactivating Cdh1. To open the M/G1 gate, Cdc20 must be activated (by successfully aligning all replicated chromosomes on the metaphase spindle); during the transition from M to G1, Cdh1 is reactivated and the G1/S gate is locked by degrading CycB. Alternation of the two switches is guaranteed by the licensing mechanism provided by the antagonism between CycB and Cdh1. Figure 3 confirms that the cartoon in Fig. 1A is indeed a precise consequence of the molecular mechanism in Fig. 1B, given reasonable assumptions on the rate laws and rate constants involved in the mathematical model.

**Figure 3.**
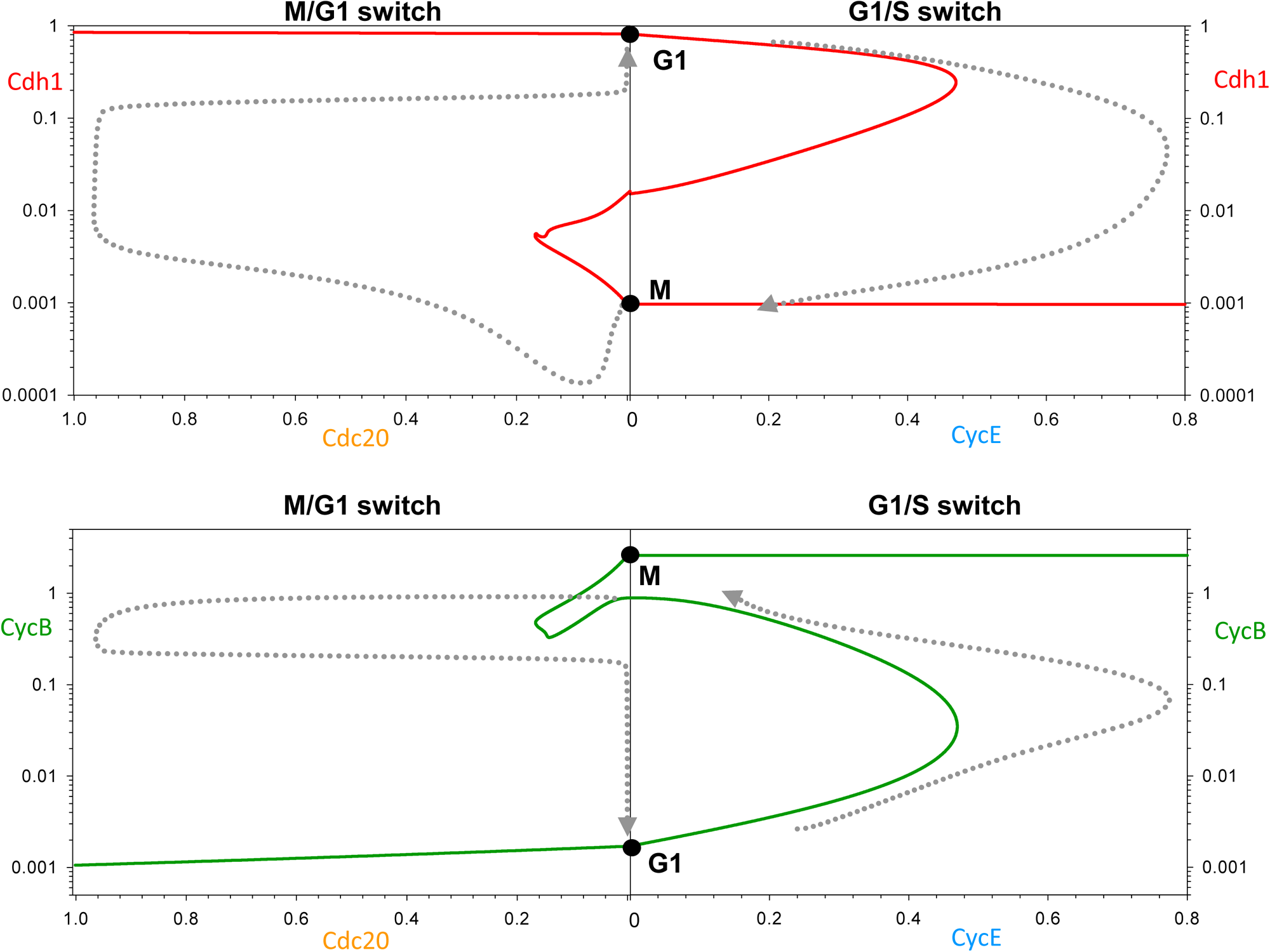
Two complementary views of progression through the mammalian cell cycle. The top (bottom) panel combines the Cdh1 (CycB) bifurcation curves in Fig. 2 (Supp. Fig. S1). In this case, the dotted lines are trajectories of the simulated limit cycle oscillation in Fig. 1. The negative feedback controls on CycE and Cdc20 are evident from the simulation, although they differ considerably from the proposed trajectories in Figs. 2 and Suppl. Fig. S1.

To provide further evidence for our model, we next discuss mutations that interfere with the alternation of the two switches.

### Endoreplication Cycles (Cdh1 Endocycles)

Mammalian cells, under certain conditions, exhibit endoreplication cycles, during which the cell undergoes multiple rounds of DNA replication without mitosis and cell division. (Under other conditions, a cell may exhibit over-replication, i.e., persistent DNA synthesis exhibiting a steady rise in DNA content.) In our view of cell cycle regulation, an endoreplicating cell does not visit the left sides of the diagrams in Fig. 3; rather it resets from G2 phase back to G1. Endoreplication can be induced in fission yeast cells by repressing synthesis of Cdc13, a B-type mitotic cyclin ^25^ and in budding yeast cells by deleting five B-type cyclins (four mitotic and one S-phase cyclin) ^26^. In fruit flies, both CycA and CycB are suppressed during endoreplication, which is driven by oscillating CycE-kinase activity ^27^. In human cells, conditional inactivation ^28^ or chemical inhibition ^29^ of Cdk1 induces discrete rounds of DNA replication without mitosis or cell division. In these endoreplicating mammalian cells, Cdh1 activity is oscillating ^29, 30^ in the absence of any Cdc20 activation; CycB level is also oscillating, although Cdk1:CycB activity is suppressed. Therefore, we classify endoreplication cycles as Cdh1**-**endocycles.

These observations are consistent with the implications of our model that the irreversible nature of the G1/S switch (under normal cell cycling) requires CycB-dependent mitotic kinase activity. To illustrate this point we have calculated the Cdh1 bifurcation diagram of the G1/S switch at different levels of Cdk1 inhibition (Fig. S2). The stronger Cdk1 inhibition is, the larger the Cdh1 reactivation threshold becomes. Above a critical value of Cdk1 inhibition (∼25% remaining Cdk1 activity), Cdh1 can reactivate at low CycE activity, rather than relying on Cdc20 activation. Observe that Cdh1 is still bistable at low Cdk1 activity (even at 0), but the G1/S switch loses its irreversible characteristic. At, say, 20% remaining Cdk1 activity, Cdh1 activity can oscillate with large amplitude as CycE activity oscillates back and forth across the two saddle-node bifurcation points (the C and Ↄ ‘noses’ of the Z-shaped bifurcation curve.

Figure 4 provides a closer view of how normal mitotic cycles are converted into Cdh1 endocycles (endoreplication cycles) as Cdk1 activity is suppressed by chemical inhibition. Mitotic cycles persist down to ∼60% inhibition of Cdk1:CycB (Fig. 4A), with the only effect to extend the duration of G2 phase (not shown). For 26-38% of remaining Cdk1 activity, our model predicts a G2 block, because

**Figure 4.**
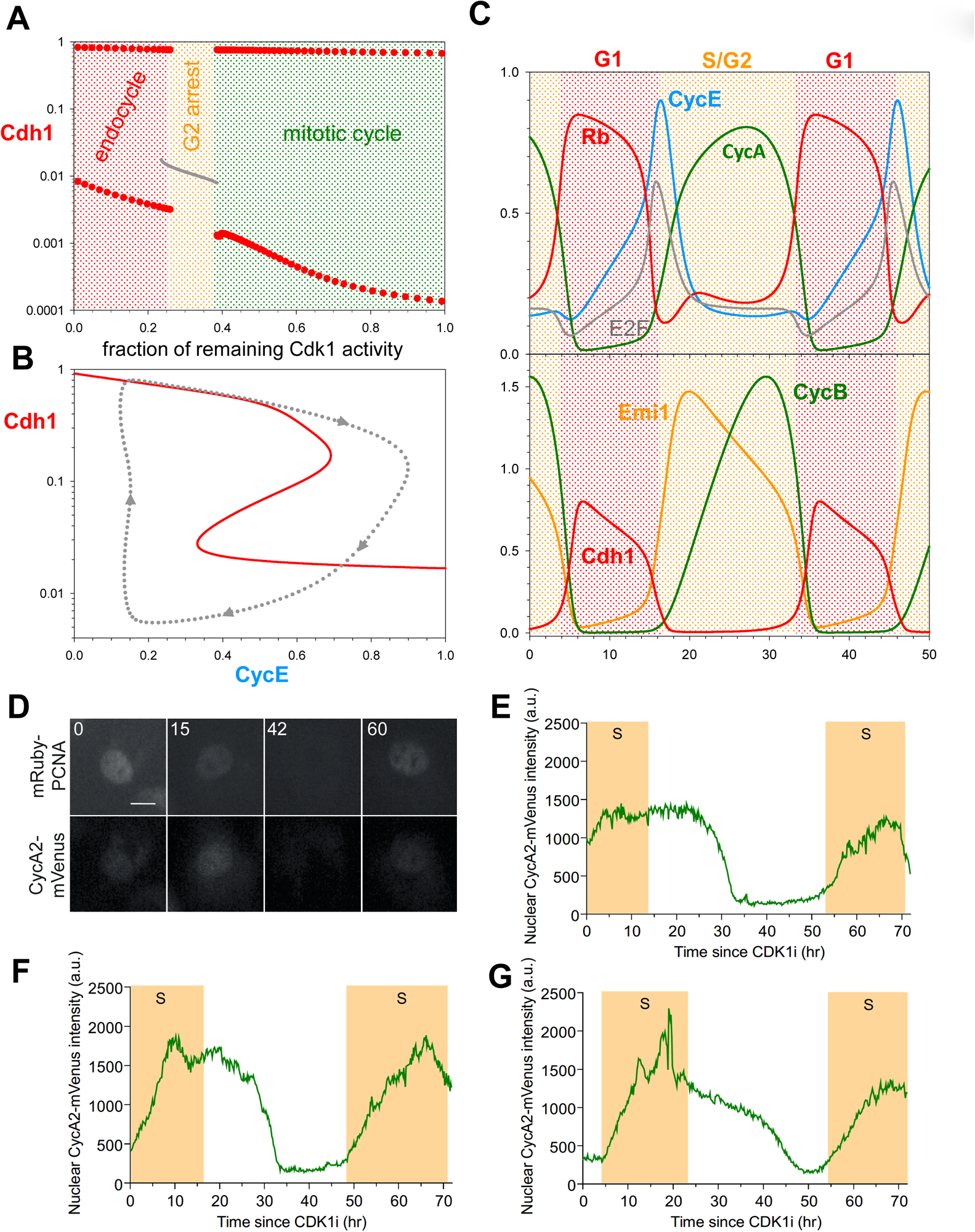
Cdk1 inhibition converts mitotic cycles into Cdh1-endocycles. (**A**) Bifurcation diagram: Cdh1 activity as a function of Cdk1 activity after chemical inhibition. Gray line: stable steady states; solid red circles: maximum and minimum excursions of Cdh1 activity during stable limit cycle oscillations. Mitotic cycles are distinguished from endoreplication cycles by the very low activity of Cdh1 (corresponding to high Cdk1:CycB activity in mitosis). (**B**) Bifurcation diagram: Cdh1 activity vs CycE, for 10% remaining Cdk1 activity. The dotted line is the projection of Cdh1 limit cycle oscillations around a hysteresis loop on the bifurcation diagram. (**C**) Simulation of Cdh1 endocycles for 10% remaining Cdk1 activity. **(D)** Still images of mRuby-PCNA and CycA2-mVenus labelled nuclei from timelapse experiments. Time shown in hours. Scale bar is 10 µm**. (E,F,G)** Graphs showing quantification of CycA2-mVenus in individual cells undergoing endocycles, plotted from the time of CDK1i addition (t = 0 h). Shaded yellow areas represent S-phase, defined by mRuby-PCNA foci.

Cdk1 is unable to self-activate by the Wee1- and Cdc25 positive feedback loops. During this G2 arrest Cdh1 is kept inactive by combined inhibition from Emi1, CycA- and CycB-kinases. Above 75% inhibition of Cdk1 activity, Cdh1 cannot be kept inactive, but rather Cdh1 executes large amplitude oscillations around a hysteresis loop involving the bistable G1/S switch only (Fig. 4B). The trajectory on the Cdh1-CycE bifurcation diagram is a projection of the simulation shown on Fig. 4C. During this limit cycle oscillation, the periodic appearance of CycE and CycA induces initiation of DNA replication, and the concomitant inactivation of Cdh1 could lead to the accumulation of the replication licensing inhibitor, geminin (not present in our model). Subsequent degradation of Emi1 reactivates Cdh1 and resets the endoreplicating cell back to G1, when replication origins can be relicensed for a new round of DNA replication. Therefore, we expect the large amplitude Cdh1 oscillations to drive discrete rounds of DNA replication characteristic of endoreplicating cells.

To experimentally test our theoretical results, we first looked for endoreplication in non-transformed hTert-RPE1 (RPE1) cells after Cdk1 inhibition with the chemical inhibitor RO-3306 (Cdk1i). After 72 h treatment with Cdk1i, we observed distinct 8n and 16n peaks by flow cytometry, indicative of endoreplication (Fig. S3A). At high concentrations of Cdk1i (>7.5 μM) an increasing fraction of cells arrest in G1 (2n), presumably due to inhibition of Cdk2 at high concentrations of RO-3306, as previously reported ^29^. In timelapse imaging using the mRuby-PCNA reporter to track DNA replication ^31^, we observed that endoreplication was even more prominent in 7.5 μM Cdk1i after depleting p53 from RPE1 cells using siRNA (Fig. S3B). Therefore, all subsequent experiments were performed under conditions of p53 depletion. To observe cell cycle dynamics in cells undergoing endocycles, we used timelapse imaging to quantify the levels of fluorescently tagged CycA2-mVenus in RPE1 cells ^32^ co-expressing mRuby-PCNA during treatment with Cdk1i. In the absence of Cdk1i, CycA-mVenus showed characteristic oscillations for mitotic cycles: peaking in intensity during cell rounding (mitotic entry) followed by abrupt degradation (Fig. S3C). In cells treated with Cdk1i, an extended G2 was observed with initially high CycA2-mVenus levels that then dropped abruptly (Fig. 4D-G and S3D,E). In 60% of these cells, this extended G2 was followed by a new round of DNA replication in the absence of any signs of mitosis (endoreplication, Fig. 4D-G and S3B). These data support our theoretical predictions.

Another way to subvert the latching gate at **M** is by suppressing Emi1 synthesis, as suggested by experiments ^33, 34^. According to our model, cells maintain their mitotic cycles up to ∼40% reduction of Emi1 synthesis (Fig. S4A). Stronger inhibition of Emi1 synthesis leads to an abrupt reduction in the amplitude of Cdh1 and Cdk1 oscillations (Fig. S4A and S4B). For nearly complete inhibition of Emi1 synthesis, the G1/S switch stops oscillating and settles onto a stable steady state. This steady state is characterized by intermediate values of Cdh1 and CycA activities, in addition to high CycE levels. We associate the reduced amplitude Cdh1 endocycles (caused by increased trough) and the intermediate Cdh1 steady states with continuous DNA synthesis (over-replication phenotype: when licensing and firing of replication origins are not temporally separated), based on the residual Cdh1 activity, which could maintain low levels of geminin, thereby allowing replication origin licensing and firing to proceed simultaneously.

### Cdc20 Endocycles

Since Cdk1 inhibition disrupts the latching property of the **M** gate and enables Cdh1 endocycles, it is tempting to consider the consequences of the opposite effect: sustained Cdk1:CycB activity. Working with HeLa cells, Pomerening *et al.* ^35^ have expressed an allele (*Cdk1AF*) for non-phosphorylatable Cdk1 subunits, which cannot be inactivated by Wee1/Myt1 inhibitory kinases. Cdk1AF short-circuits the Cdk1 activation feedback loop operating at the G2/M transition (Fig. 1B). Cdk1AF-expressing cells carry out a relatively normal first mitosis, but then undergo rapid cycles of CycB accumulation and degradation at 3-6 h intervals. These fast CycB oscillations show certain resemblances to the early embryonic cell cycles of Xenopus. Inspired by these experimental results, we decided to analyze the effects of weakening inhibitory Cdk1 phosphorylation in our model (Fig. 5).

**Figure 5.**
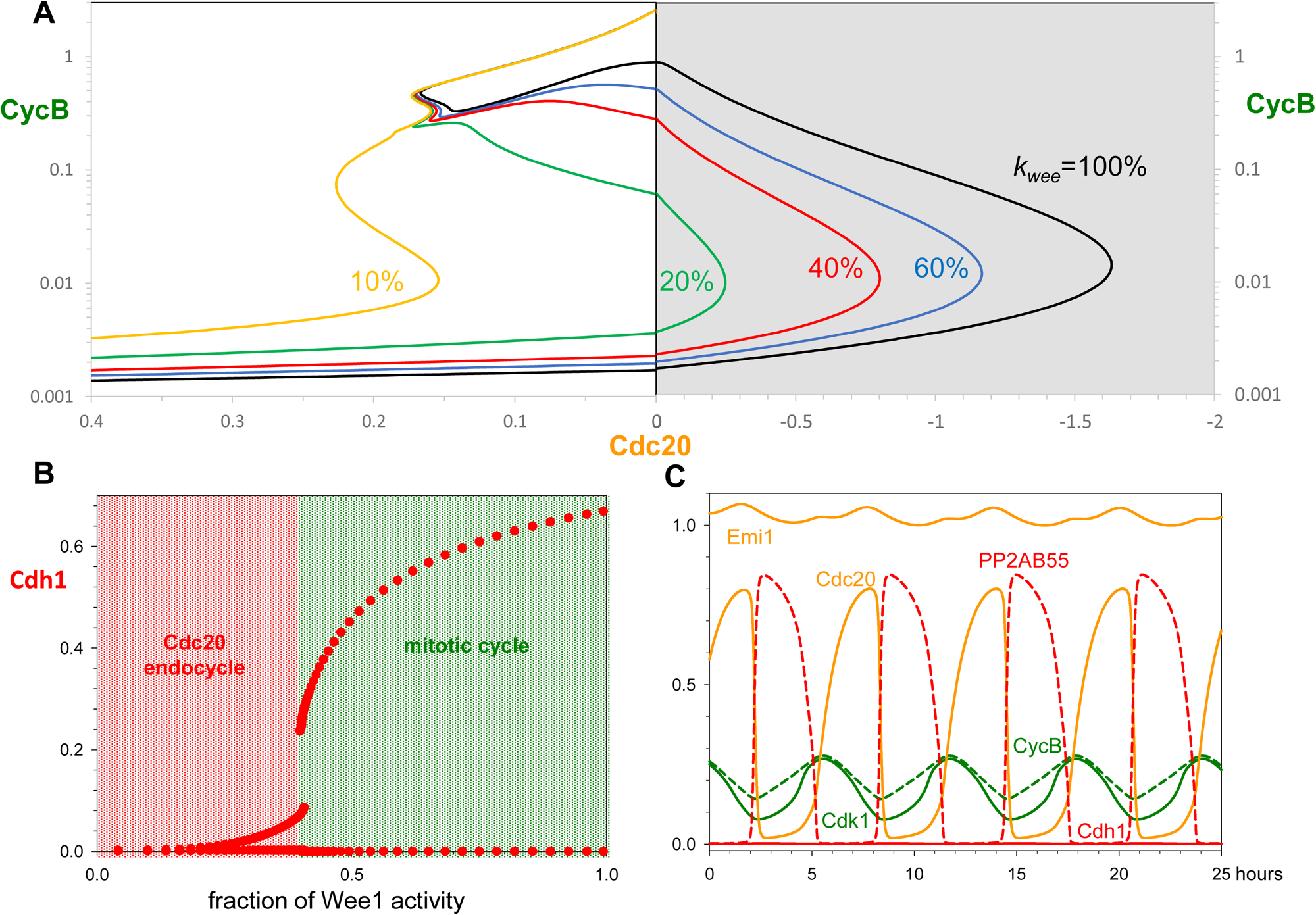
Inhibition of Wee1 kinase activity converts mitotic cycles into Cdc20 endocycles. (**A**) Bifurcation diagram: CycB activity as a function of Cdc20, for increasing inhibition of Wee1. (**B**) Bifurcation diagram: Cdh1 activity as a function of remaining Wee1 activity. (**C**) Simulated Cdc20 endocycles, for 10% Wee1 activity.

Figure 5A presents CycB vs Cdc20 bifurcation diagrams for different values of Wee1 activity (*k_wee_*). Decreasing Wee1 activity moves the threshold for Cdc20 inactivation (the threshold for CycB re-accumulation) to less negative values of Cdc20 (i.e., to the left in Fig. 5A). When Wee1 activity falls below 13%, the Cdc20 threshold for CycB re-accumulation moves to positive values of Cdc20, meaning that exit from mitosis no longer latches the cell at the **G1** gate. Now the control system can oscillate around a hysteresis loop on the CycB-Cdc20 bifurcation diagram. As the inhibitory phosphorylation of Cdk1 becomes weaker, the amplitude of the Cdh1 oscillations decreases (Fig. 5B) and finally becomes negligible below 25% of Wee1 activity. In the absence of any fluctuations of Cdh1, the CycB level still shows persistent oscillations at low Wee1 activity (Fig. 5C). These oscillations of CycB level are exclusively driven by fluctuating activity of APC/C:Cdc20; so we call them Cdc20 endocycles. During Cdc20 endocycles, Cdh1 is kept inactive by high Emi1 levels and by strong inhibition by Cdk1:CycB kinase (Fig. 5C). Since the synthesis of both Cdh1 inhibitors is dependent on E2F activity (directly for Emi1 and indirectly—via CycA—for CycB), sustained Cdc20 endocycles require that the level of Rb must be less than the level of E2F. Indeed, these limit cycle oscillations persist in the absence of Rb, providing an explanation for the observations of Pomerening *et al.* ^35^ of Cdc20 endocycles in Rb-negative HeLa cells. We have experimentally tested for Cdc20 endocycles in Rb-positive RPE1 cells, which will be discussed after describing Rb’s role in the cell-size checkpoint.

In summary, we have shown that inhibition and premature activation of the mitotic kinase has opposite effects on human cell-cycle switches. Cdk1 inhibition breaks the latch at the M/G1 gate and induces Cdh1 endocycles, which trigger periodic and distinct rounds of DNA replications. In contrast, in the absence of inhibitory Cdk1 phosphorylation, the G1/S latch is broken, and CycB level oscillates rapidly by the periodic activation/inactivation of Cdc20.

### Checkpoints

Up to this point we have been treating the cell-cycle control network as an oscillator, which induces cell cycle events by measuring time only. However, this underlying clock is subject to several checkpoint mechanisms that make progression through the cell cycle sensitive to a variety of important intra- and extracellular signals. The most important signals are (1) extracellular growth- and antigrowth factors, which govern passage through the restriction point, (2) cell growth, which must be sufficient to authorize the G1/S transition, (3) DNA damage, which can block both G1/S and the G2/M transitions, (4) unreplicated DNA, which blocks mitotic entry, and (5) misaligned chromosomes, which prevent the metaphase-to-anaphase transition. These checkpoint mechanisms stop progression around the cell-cycle loop (Fig. 3) by creating stable steady states on the upper and lower branches of the bifurcation curves near the neutral point, where both CycE and Cdc20 are absent. In this subsection we focus on two exemplary checkpoints.

The mitotic checkpoint blocks activation of Cdc20 (thereby inhibiting degradation of CycB and securin) until all chromosomes become bioriented on the mitotic spindle ^3^. (Upon degradation of securin, active separase cleaves the cohesin rings that are holding sister chromatids together at bioriented centromeres, allowing the sister chromatids to be separated in anaphase.) In the model, a reduction of Cdc20 activity below about 10% normal (not shown) terminates the limit cycle oscillation of CycB and creates a stable steady state of Cdk1:CycB activity.

The effects of cell growth on cell cycle progression are complex and as yet not fully understood. However, it has been demonstrated that Rb plays an important role in size control ^36^. Above a certain threshold concentration, Rb inhibits the G1/S transition by blocking E2F-dependent expression of CycE, CycA and Emi1. Our model is consistent with this observation because, at high Rb concentration, large amplitude mitotic oscillations of CycB become stabilized at a low, steady state concentration, characteristic of G1 phase (Fig. S5A). To illustrate the role of Rb in cell-size control, we have supplemented our clock mechanism with an Rb-dilution model ^36^. We assume that cells are growing linearly in volume and that the Rb synthesis rate is size-independent (proportional to the genome content) and transcriptionally regulated. Fast Rb synthesis is restricted to a four-hour long window starting around the G1/S transition and leading to a doubling of Rb concentration; subsequently, Rb concentration is diluted out by volume growth during the remainder of the cycle (Fig. 6). Our assumptions provide a temporal pattern for cell cycle changes in the amount of Rb molecules (Fig. S5B) that agrees well with the experimental data of Zatulovskiy *et al.* ^36^. In this framework, Rb concentration (amount/volume) mirrors the cellular DNA/volume ratio and provides a possible mechanism for balanced growth and division, by adjusting the period of the cell cycle to the time required to double cell mass (see Fig. S5A).

**Figure 6.**
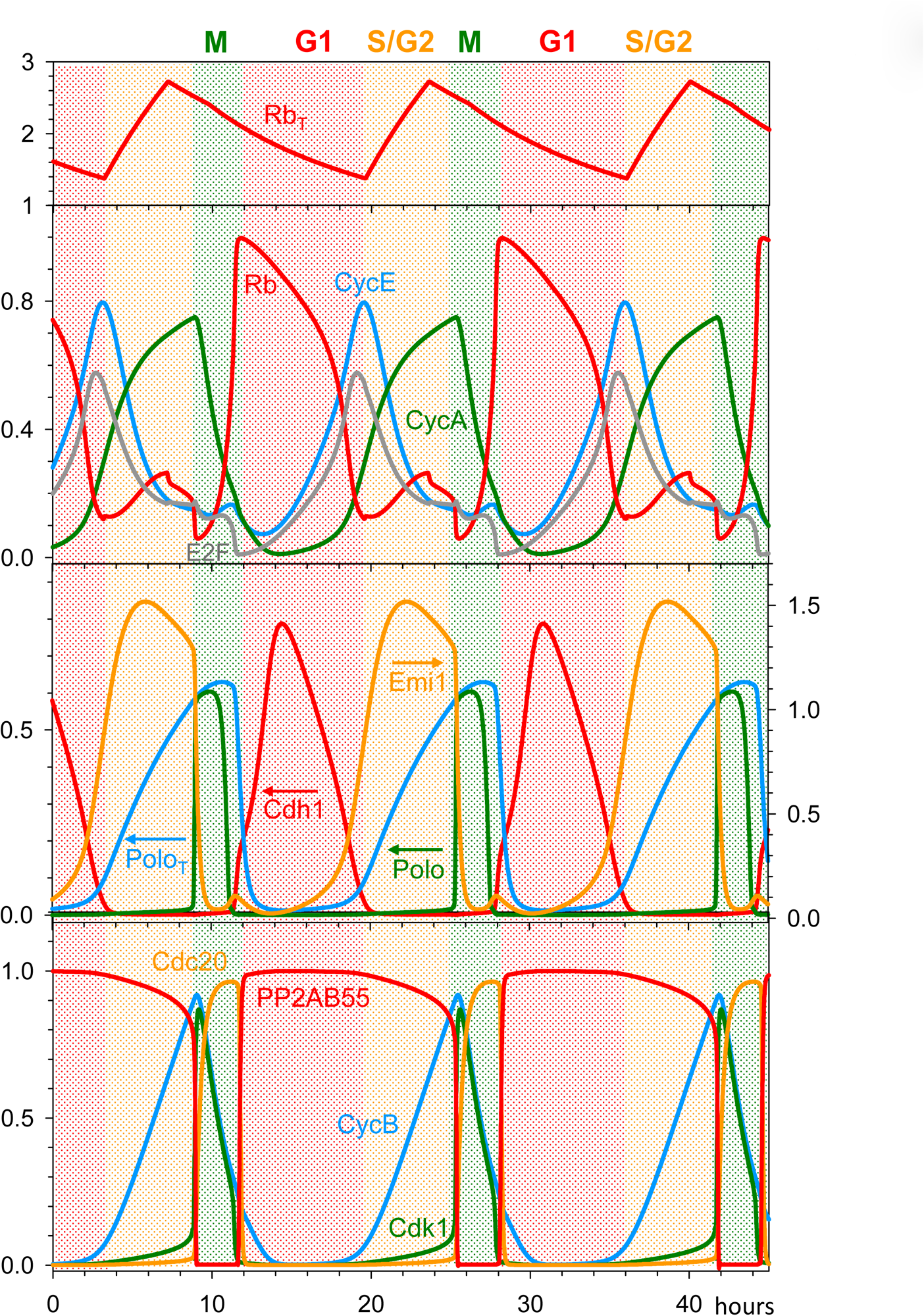
Growth-controlled cell cycle by Rb-dilution. The limit-cycle model is supplemented with cell cycle-regulated transcriptional control over Rb synthesis. Rb synthesis during S/G2 phase results in an increase of its concentration, followed during the remainder of the cell cycle by decreasing Rb concentration due to dilution by cell volume growth. (Notice that Rb concentration does not change during cell division.) This mechanism automatically leads to two-fold fluctuations in Rb concentration when cell volume doubles over the course of a cell cycle.

### Rb-controlled Cdc20 Endocycles

We have tested the possibility that constitutively active Cdk1:CycB could induce Cdc20 endocycles in the context of size control by an Rb-dilution mechanism. Our model predicts that inactivation of Wee1 after completion of mitosis induces small amplitude oscillations in CycB level, while Cdh1 is completely inhibited (Fig. 7A). Moreover, these Cdc20 endocycles have a period very close to the normal cycle time, because they are controlled by periodic synthesis and dilution of Rb in the following way. Cdc20 endocycles are driven by the fundamental negative feedback loop between CycB and Cdc20 (Cdk1:CycB activates APC/C:Cdc20 and APC/C:Cdc20 degrades CycB). Since CycB synthesis is initiated by CycA-dependent kinase and CycA is synthesized by E2F transcription factor in an Rb-dependent manner, Cdc20 endocycles (in Rb-positive cells) are controlled in part by the oscillating level of unphosphorylated Rb. Whenever unphosphorylated Rb is in stoichiometric excess over E2F the synthesis of both CycA and CycB are on hold and the oscillation is temporarily stopped.

**Figure 7.**
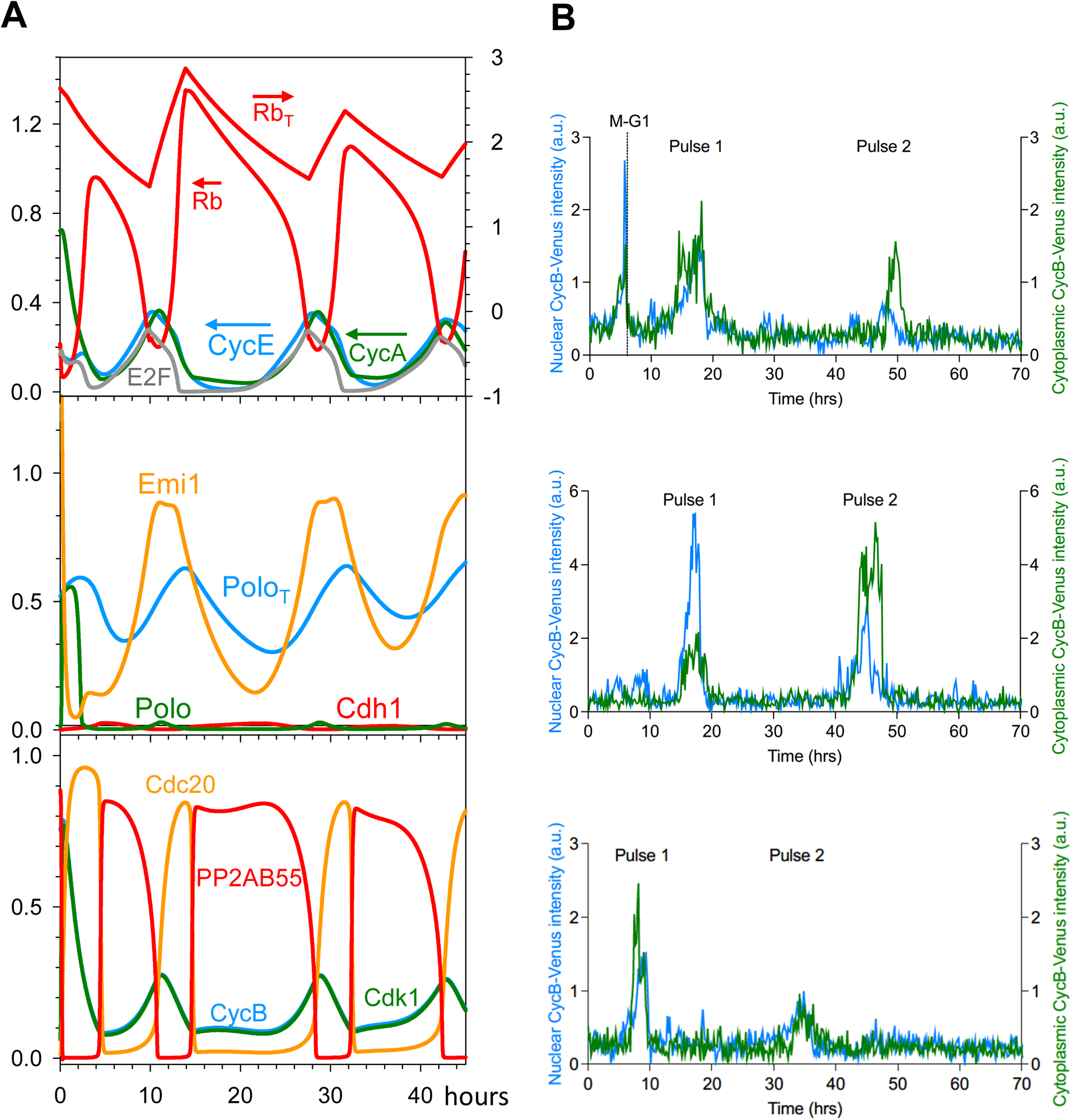
Cdc20 endocycles controlled by Rb-dilution. **(A)** Numerical simulation of the growth controlled cell cycle model with Wee1 inhibition (k_wee_=0). After exiting mitosis, CycB level shows small amplitude oscillations driven by APC/C:Cdc20 in the absence of any Cdh1 activity. Since Cdk1:CycB activity does not reach the mitotic threshold, both nuclear and cell division are hampered. The continuous rise in cell volume (not shown) causes an imbalance with Rb synthesis and it creates a decreasing amplitude in concentration of Rb oscillations. (**B)** Normalised CycB1-mVenus intensity in individual cells treated with Wee1 siRNA and undergoing Cdc20 endocycles. Blue curve: nuclear CycB level; green curve: cytoplasmic CycB level.

We have used siRNA to deplete Wee1 inhibitory kinase in order to induce constitutively active Cdk1:CycB complexes in Rb-positive RPE1 cells. In RPE1 cells with fluorescently-tagged CycB1-mVenus ^37^, we used timelapse imaging to quantify CycB1 protein levels after Wee1 depletion. In control-depleted cells, CycB1-mVenus oscillates – increasing prior to mitotic entry (defined by cell rounding) and rapidly degraded upon mitotic exit (Fig. S6A). After Wee1 depletion by siRNA (Fig. S6B), cells may go through an initial early mitosis but then cells continue to grow in volume, becoming large, interphase-arrested cells. Despite their robust interphase arrest, we observed cells that displayed one or two bursts of CycB signal, both in the cytoplasm and in the nucleus (Fig. 7B; S6C,D). The rise in CycB level was not accompanied by nuclear division. This supports our theoretical prediction that in the absence of Cdk1 inhibitory phosphorylation, cells are undergoing Cdc20 endocycles.

In summary, our results support and supplement the findings of Pomerening *et al.* ^35^, who first described small amplitude CycB oscillations in the absence of Cdk1-inhibitory phosphorylation in HeLa cells. In Rb-negative HeLa cells, Cdc20 endocycles behave as an autonomous oscillator ^35^, while Rb-positive RPE1 cells, the period of Cdc20 oscillations is influenced by an Rb-mediated size-control mechanism (our work).

## DISCUSSION

We have previously proposed that G1 and M are two alternative stable steady states of the budding yeast cell cycle control system ^5–7^. These alternative steady states are the consequence of double-negative feedback between B-type (Clb1-5) cyclin-dependent kinases (B-CDKs) and their antagonists (APC/C:Cdh1 and Sic1, a stoichiometric CDK inhibitor). Our toggle-switch concept of the yeast cell cycle has been verified by elegant experiments in budding yeast ^38, 39^. Recently, we have shown that the toggle-switch model also provides a natural explanation for different endocycles induced by perturbation of mitotic cyclin expression ^7^:

i. Endoreplication: discrete rounds of DNA replication induced by deletion of Clb1-4 (the mitotic cyclins) and of Clb5 (one of the S phase cyclins) ^40^.
ii. Cdc14 endocycles: periodic activation of the Cdc14 mitotic-exit phosphatase in the presence of non-degradable mitotic cyclin, Clb2 ^41, 42^.

In yeast, Cdh1 activity oscillates during both endocycles, and it promotes the degradation of the Nrm1 transcription inhibitor and of polo-kinase (Cdc5) during endoreplication and Cdc14 endocycles, respectively.

The mutual antagonism between the protein degradation pathway initiated by APC/C:Cdh1 and its target proteins CycA, CycB and Emi1 suggests that our toggle-switch concept also applies to the mammalian cell cycle. Indeed, hysteresis in the regulation of APC/C:Cdh1 activity is supported by experiments with mammalian cells ^13^. Our hypothesis is illustrated schematically in Fig. 1A. The bistable toggle switch (between APC:Cdh1 and Cdk1:CycB) is flipped ‘on’ (high CycB:Cdk1 activity) by Cdk2:CycE and flipped ‘off’ (high APC:Cdh1 activity) by APC:Cdc20. We find that inhibition of mitotic CycB:Cdk1 complex makes APC/C:Cdc20 dispensable for Cdh1 reactivation by disabling the ‘latching’ property mitotic steady state (**M**), converting the ‘one-way’ toggle switch into an autonomous oscillator regulated only by the remaining antagonistic interactions between APC/C:Cdh1 and CycA:Cdk2 + Emi1. In the absence of mitotic CDK activity, cells are driven around a Cdh1-hysteresis loop by negative feedback regulation of CycE-kinase activity. The oscillations in CycE and CycA levels and their CDK activities lead to discrete rounds of DNA synthesis, analogous to yeast endoreplication cycles. We have confirmed this by live-cell imaging of fluorescently tagged CycA in RPE1 cells exposed to a Cdk1 inhibitor, RO3306.

This mechanism of endoreplication, suggested by our theoretical model and verified experimentally, provides a basis for understanding how whole genome doubling (WGD) can arise during tumorigenesis. The many layers of regulation underlying our ‘latching’ mechanism for cell cycle progression, ensure that WGD is a rare event. However, it is estimated that up to 40% of all cancers have undergone at least one WGD event ^43^. WGD can promote tumorigenesis by buffering the effects of deleterious mutations, by fostering mutations that increase cell proliferation ^44–46^, and— quite generally—by disrupting the genomic stability of cells ^47^. By providing a mechanistic basis for how WGD can arise, our model might assist efforts to develop targeted treatments against WGD.

Inducing mitosis in the presence of non-degradable CycB represents the inverse perturbation of mitotic CDKs, corresponding approximately to Cdc14 endocycles in yeast cells. In mammalian cells, persistent mitotic Cdk1 activity induced by non-degradable CycB reactivates the error-correction mechanism of the mitotic checkpoint, which results in oscillating sister-chromatids between the two poles (pseudo-anaphase) ^48, 49^. These oscillations are the consequence of tension-dependent fluctuations of Aurora-B kinase activity at kinetochores. We have investigated an alternative way to disrupt the antagonistic relationship between mitotic kinase and APC/C:Cdh1 in mammalian cells, by depleting cells of Wee1 kinase, the kinase that inhibits CycB:Cdk1 activity in G2. We have found that sustained activity of CycB:Cdk1 in Wee1-depleted cells makes CycE-kinase dispensable for Cdh1-inactivation, because it maintains Cdh1 constitutively phosphorylated and inactive. Moreover, in the absence of inhibitory phosphorylation of CycB:Cdk1, APC/C:Cdc20 is activated prematurely, which promotes early degradation of CycB and (because of the negative feedback loop between CycB and Cdc20) loss of APC/C:Cdc20 activity. Hence, although CycB:Cdk1 activity is ‘sustained’ under these conditions, the amplitude of CycB:Cdk1 oscillations is never high enough to drive the cell into mitosis or low enough to let Cdh1 make a come-back. Therefore, sustained activity of CycB:Cdk1 induces Cdc20 endocycles in the absence of Cdh1 activity, which makes the situation in human cells different from yeast’s Cdc14 endocycles where Cdh1 oscillates. This dissimilarity between yeast and human cells could be a consequence of different mitotic exit phosphatases and their regulation, as well as different roles of Cdc20 and Cdh1 in the degradation of mitotic CycBs. In budding yeast, complete degradation of Clb2 mitotic cyclin requires Cdh1, which is dephosphorylated during mitotic progression by the release of active Cdc14 phosphatase from the nucleolus ^50^. In contrast, in human cells Cdh1 is dispensable for degradation of mitotic cyclins, and their mitotic exit phosphatase, PP2A:B55, is kept inactive by CycB:Cdk1 via the Greatwall-ENSA pathway. Despite this differences, notice that Cdc20 fluctuations induced by sustained CycB:Cdk1 activity are accompanied by large amplitude oscillations of PP2A:B55 phosphatase activity (Fig. 7A). This observation suggests that unregulated Cdk1 activity induces mitotic exit phosphatase endocycles in both yeast and human cells.

On the experimental front, we have demonstrated these small amplitude oscillations in CycB levels using live-cell imaging of RPE1 cells depleted for Wee1 by siRNA. The period of the oscillations were significantly longer than the CycB oscillations observed by Pomerening et al. ^35^ in HeLa cells, which we can explain by the indirect role of an Rb-dependent size-control mechanism in the transcription of CycB. Significantly, Wee1 inhibitors are currently in clinical trials for cancer treatment. The aim of these inhibitors is to specifically target cancer cells on the basis that only p53-mutant cancers, which rely on Wee1 to maintain the DNA damage checkpoint in G2, will be sensitive to Wee1 inhibitors ^51, 52^. By providing an understanding of the effects of inhibiting Wee1 in non-cancerous cells, our model may allow for a better understanding of potential side-effects.

We have simplified our human cell cycle model by neglecting some cell cycle regulators, including cyclin-dependent kinase inhibitors (CKIs) like p27, p21 etc. These CKIs provide an extra layer of antagonism to the regulatory network (CKIs inhibit CDKs and are targeted to degradation by CDKs). There is no theoretical bottleneck to expand our model with CKIs, and this is a task for future work. For instance, p27 has a complex role in regulating the activities of CycD:Cdk4/6 and CycE:Cdk2 ^53^, thereby influencing the G1/S transition by interfering with the Rb-E2F double-negative feedback loop. p21 plays similar roles in the DNA-damage response induced by p53.

## Materials and methods

### Computational methods

#### Cell cycle clock model

The mathematical model presented here describes the biochemical interactions governing the mammalian cell cycle control network. It is assumed that the activity of Cyclin:CDK complexes is limited by the availability of cyclins, which strongly and rapidly bind CDKs. In early G1, cyclin expression is repressed via Rb-dependent stoichiometric inhibition of E2F transcription factors. A fraction of total Rb protein is mono-phosphorylated and inactivated by CycD:CDK4/6 (a parameter, here); the remaining fraction of unphosphorylated Rb is:

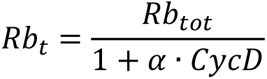

This pool of unphosphorylated Rb can be further phosphorylated by the other Cyclin:CDK complexes, such that the rate law of Rb available to inhibit E2F (i.e., Rb molecules that are unphosphorylated by any Cyclin:CDK complexes) is given by the differential equation:

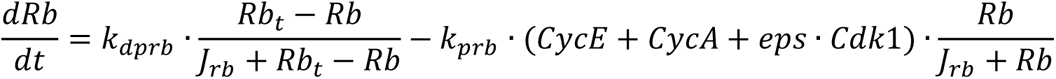

We are using Michaelis-Menten kinetics to describe the rates of phosphorylation (‘prb’) and dephosphorylation (‘dprb’) of Rb. Next, assuming that the Rb:E2F complex (RbE2F) is in equilibrium with the dissociated monomers, we calculate its concentration by:

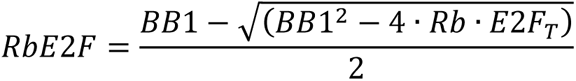

where *BB*1 = *Rb* + *E*2*F*_*T*_ + *K*_*drbe*2*f*_, *E2F_T_* is the total concentration of E2F (assumed to be constant), and *K_drbe2f_* is the equilibrium-dissociation constant of the complex. In addition, E2F can be independently inhibited through CDK-dependent phosphorylation:

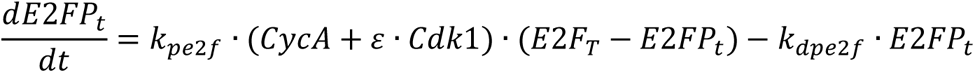

Consequently, the fraction of active E2F is given by:

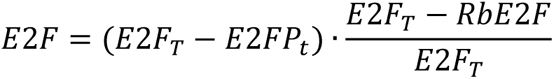

Active E2F (i.e., unbound by Rb and unphosphorylated by CDKs) stimulates the transcription of a number of genes required for G1/S progression, including CycE, CycA and Emi1.

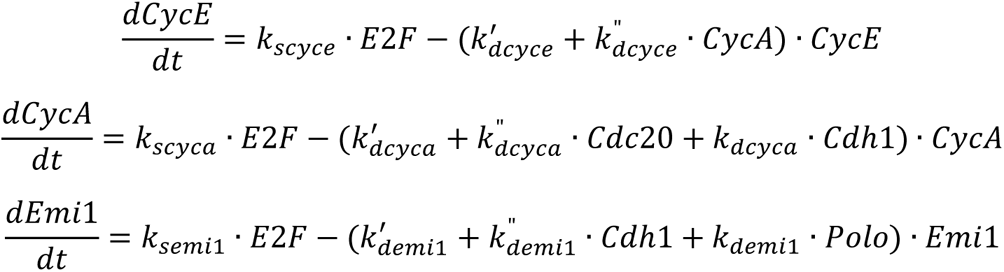

In addition to being regulated transcriptionally, these proteins are also targeted for degradation in specific manners, as described by the ‘*k_d…_*’ terms in these differential equations. CycA is a substrate of the ubiquitin ligase APC/C in complex with either Cdc20 or Cdh1; CycE is a substrate of the SCF ubiquitin ligase, after it is phosphorylated by CycA:Cdk2; and Emi1 is a target of both APC/C:Cdh1-mediated degradation and SCF-mediated degradation (after phosphorylation by Polo kinase). For these reasons, CycE—but not CycA or Emi1—accumulates in G1; in S phase, CycE is rapidly degraded in response to CycA-mediated phosphorylation; and during M phase, both CycA and Emi1 are rapidly degraded (by different pathways) and kept low throughout G1. All of these regulators cooperate to drive the inactivation of Cdh1 at the G1/S transition. The stoichiometric binding of Emi1 to Cdh1 is modelled in the same way as the binding of Rb to E2F, namely:

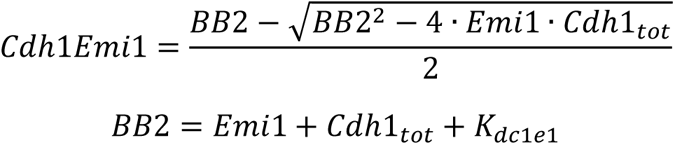

The phosphorylation of Cdh1 by CycE, CycA and CycB is described by:

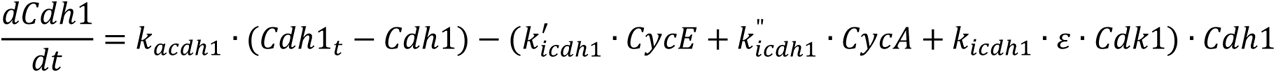

where *Cdh1_t_* is the Emi1-free Cdh1: *Cdh*1_*t*_ = *Cdh*1_*tot*_ − *Cdh*1*Emi*1.

As CycA accumulates, it is responsible for driving the accumulation of CycB and Polo:

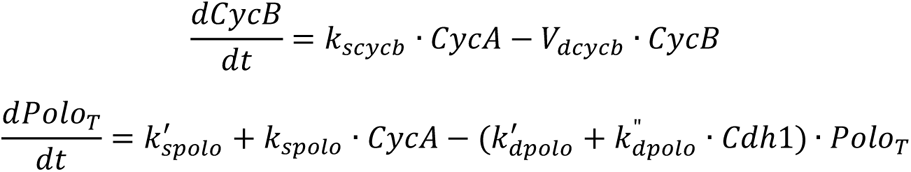

with *V_dcycb_* being a degradation rate function that depends on Cdc20 and Cdh1:

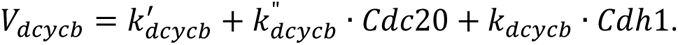

Nevertheless, as the CycB:Cdk1 complex accumulates, it is initially inactivated by Wee1-dependent phosphorylation; the active, dephosphorylated form is denoted as Cdk1:

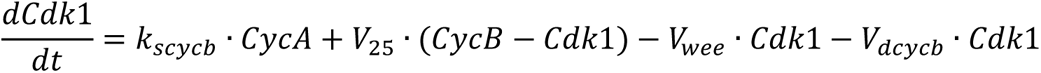

The net rate of accumulation of the dephosphorylated CycB:Cdk1 complex depends on the rate functions for the Wee1 kinase and Cdc25 phosphatase reactions:

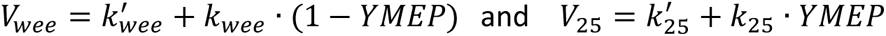

where YMEP is a Goldbeter-Koshland function for the tyrosine-modifying enzymes:

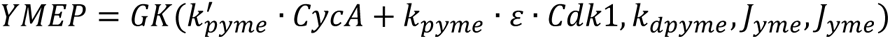

The GK function depends on the activities of CycA, Cdk1 and a constitutive phosphatase, denoted by the constant parameter *k_dpyme_*. The GK function is defined as:

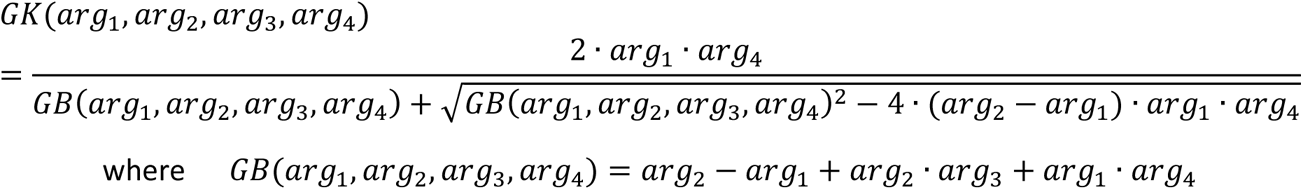

The GK-function describes the steady state phosphorylated/dephosphorylated ratio of a substrate that is subject to a kinase-phosphatase couple operating near saturation. The response is sigmoidal with respect to the kinase/phosphatase activity ratio.

Together, CycA and Cdk1 also lead to the activation of Polo and Greatwall kinases:

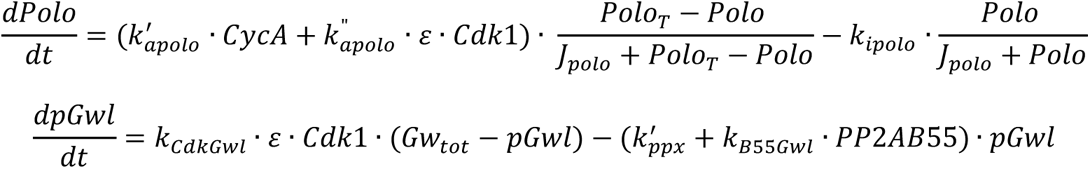

Notably, *ε* is a parameter that quantifies the relative activity of Cdk1. We set *ε* =1, unless it is reduced to a value 0 < *ε* < 1, to simulate Cdk1 inhibition, as indicated in the text. In addition, Gwl is dephosphorylated by the PP2A:B55 phosphatase. In its active, phosphorylated form, Gwl phosphorylates ENSA (pENSAt), which leads to the formation of an inhibitory complex with the PP2A:B55 phosphatase.

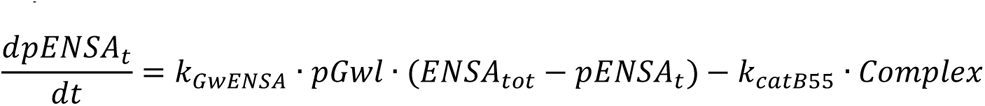

Where *Complex* = *B*55_*tot*_ − *PP*2*AB*55, *B55_tot_* being the total concentration of B55, assumed to be constant. The dissociation of the complex is favoured by the PP2A:B55-dependent dephosphorylation of pENSA:

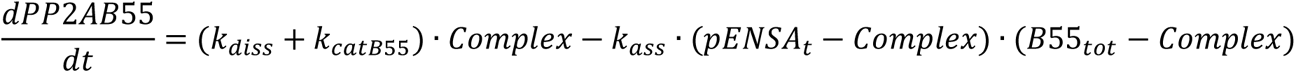

Finally, when the ratio of Cdk1 and PP2AB55 increases sufficiently, Cdc20 is activated, leading to the degradation of mitotic cyclins:

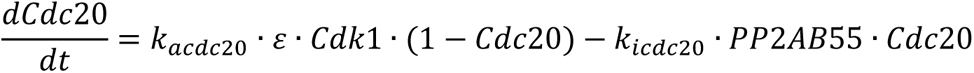

#### Size control model

The rate of cell volume growth is assumed to be constant (see Fig. 1K in ^36^):

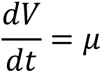

and volume is halved at cell division when Cdk1 drops below 0.7. In order to model size-controlled cycling, the total Rb concentration is converted from a constant to a dynamic variable, where the rate of synthesis in concentration units is inversely proportional to the volume:

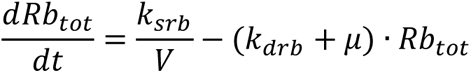

The rate of Rb synthesis (*k_srb_*) is assumed to change in a cell cycle dependent manner. During G1, *k_srb_* is very small (0.02h^-1^), which means that the amount of total Rb protein is roughly constant, given a sufficiently long (∼ 30h) half-life (*k_drb_* = 0.023 h^-1^). Consequently, the protein concentration depends on the cellular volume at this stage, or in other words, the rate of change of Rb_tot_ concentration depends on the rate of volume growth, *µ*. Nevertheless, the amount of Rb must be replenished during each cycle; to this end, we assume that Rb expression is turned on (*k_srb_* = 0.1 h^-1^) after S-phase entry (when CycA > 0.3) for a fixed duration (4 h), ensuring that a fixed amount of protein is expressed during each cycle. This amount corresponds to a doubling of the Rb number of molecules present in early G1.

### Computation

Solutions to the system of differential equations introduced above have been calculated numerically, using the XPPAUT software package with the ‘Stiff’ integration method. The numerical values of the parameters are provided in Table 1, unless otherwise stated.

### Bifurcation diagram calculation

Bifurcation diagrams of the system were calculated using the AUTO extension of XPP. Given our assumption that there is no significant activity overlap between the two helper molecules, CycE and Cdc20, the differential equations describing the two species were replaced by parameters with the same name. Thus, to plot the bifurcation diagrams with respect to CycE, Cdc20 was set to zero, and the steady state solutions of the system were calculated for a range of CycE values. Cdc20 bifurcation diagrams were calculated analogously.

### XPPAUT code for simulation of the mammalian cell cycle model

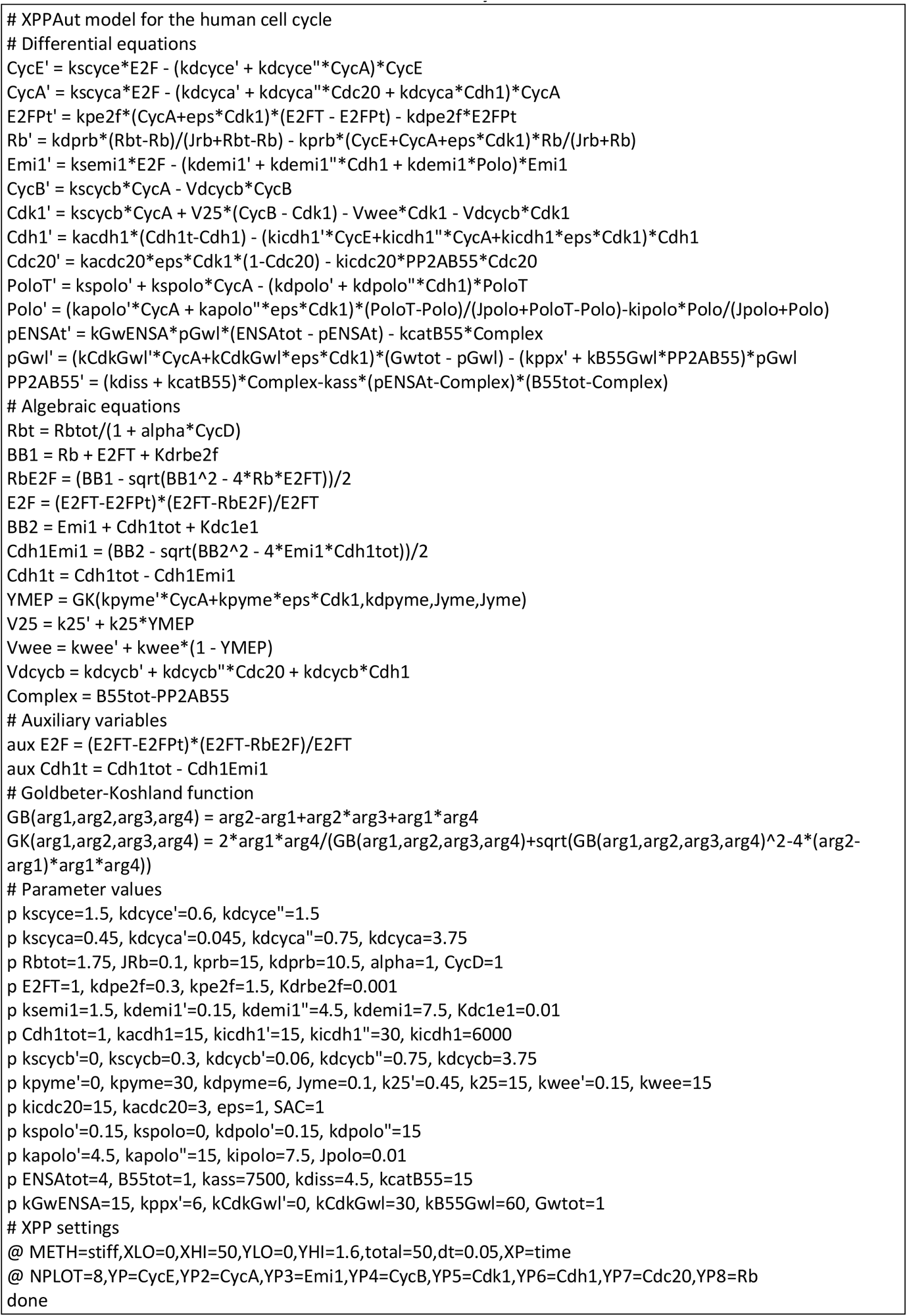

### Experimental methods

#### Cell maintenance

hTert-RPE1 cells were maintained in DMEM with 10% FBS and 1% P/S at 37°C and 5% CO_2_. Cells were passaged every 3-4 days and tested for mycoplasma by ELISA every month.

#### Cell cycle analysis by Flow Cytometry

hTert-RPE1 cells were seeded at 30% confluency into 6 well tissue culture plates one day before treatment. The next day, DMSO (vehicle control) or different concentrations of the CDK1i, RO-3306, were added to wells and left in for 72 h. After 72 h, cells were washed 1X in PBS, trypsinised and centrifuged at 1000xg for 5 min at 4°C. The cell pellet was washed one more time in PBS, before cells were resuspended in 300 ul of PBS. Cells were then fixed by adding 700 ul of 100% ice-cold ethanol at kept at -20°C overnight. The next day, cells were washed in ice-cold PBS and stained with 20 μg/ml PI solution in PBS/0.1% TritonX-100 and 200 μg/ml DNAse-free RNAseA (ThermoFisher) for 30 mins, in the dark at RT. Stained cells were strained through 0.2 μm cell strainers into FACS tubes (BD) and analysed on the BD FACS Symphony analyser. Cell cycle analysis was performed in FlowJo.

#### Cyclin A2-mVenus timelapse experiments

hTert-RPE1 mTurquoise-H2B mRuby-PCNA Cyclin A2-Venus cells were reverse transfected in 384w PhenoPlates (PerkinElmer) with non-targeting control (NTC) or TP53 siRNA (ONTargetpools, Horizon Discovery). Cells were transfected with 20 nM siRNA using 40 nl/well of Lipofectamine RNAiMAX (FisherScientific) diluted in 10ul OPTIMEM. 1000 cells/well were plated on top of the transfection mix in 20ul of phenol-red free DMEM (with 10% FBS and 1% P/S) and incubated for 24h. Before imaging, cells were treated with either DMSO or 7.5 µM of the CDK1 inhibitor (CDK1i), RO-3306. A breathable membrane was applied over the plate and cells were imaged every 10 min on the Operetta CLS high-content microscope (PerkinElmer) at 37°C and 5% CO_2_ for 72h using the 20x N.A. 0.8 objective. Background subtraction was performed in FIJI and cells were manually tracked to determine S-phase and quantify CycA2-mVenus intensity.

#### Cyclin B1-mVenus timelapse experiments

hTert-RPE1 Cyclin B1-mVenus cells were reverse transfected in Ibitreat 8-well Ibidi chambers with NTC (non-targeting control) or Wee1 siRNA (ONTargetpools, Horizon Discovery). Cells were transfected with 20 nM siRNA using 0.16 ul Lipofectamine RNAiMAX (FisherScientific) diluted in 40 ul/well OPTIMEM. 6000 cells/well were plated on top of the transfection mix in 300ul of phenol-red free DMEM (with 10% FBS and 1% P/S) in and left for 6h. Cells were imaged every 10 mins on the inverted Olympus IX83 microscope with spinning disk unit at 37°C and 5% CO_2_ for 72h using the 20x N.A. 0.7 objective. Background subtraction was performed in FIJI and cells were manually tracked to quantify CycB1-mVenus nuclear intensity.

#### Western blotting

hTert-RPE1 Cyclin B1-mVenus cells were reverse transfected in 24 well plates with 20 nM NTC or Wee1 siRNA using 1ul of Lipofectamine RNAiMAX (FisherScientific) diluted in 100 ul/well OPTIMEM. Cells were plated on top of the transfection mix in 400ul of DMEM (with 10% FBS and 1% P/S) and left for 6h. Cells were then washed in 1X PBS lysed in 50ul of 1X Laemmli buffer and cell lysates were loaded and run into 4-20% Tris-Glycine Novex pre-cast gels (FisherScientific). Separated proteins were transferred to PVDF-FL membranes which were then blocked in blocking buffer (5% milk in TBS with 10% glycerol) for 1hr at RT. Antibodies raised against Wee1 (CST 4936, 1:500) and Vinculin (CST 13901, 1:2000) were diluted in blocking buffer and incubated with membranes overnight at 4°C. The next day, membranes were washed 3 x 10 mins in TBS/T (TBS with 0.05% TritonX-100) and then incubated for 1hr at RT in anti-rabbit secondary antibody conjugated to HRP diluted 1:2000 in blocking buffer. Membranes were washed 3x 10 mins in TBS/T and developed using Biorad ECL substrate. Blots were imaged on an Amersham Imager 600.

## Acknowledgements

We acknowledge financial support from BBSRC Strategic LoLa grant BB/M00354X/1 to BN. We would like to thank Jonathon Pines (ICR, London) for sharing the hTert-RPE1 Cyclin B1-mVenus cell line and Joerg Mansfeld (ICR, London) for sharing the hTert-RPE1 mTurquoise-H2B mRuby-PCNA Cyclin A2-Venus cell line. ARB is funded by a CRUK Career Development Fellowship (C63833/A25729) and her lab receives core-funding from the MRC-LMS (MC-A658-5TY60). We thank the MRC-LMS/NIHR Imperial Biomedical Research Centre Flow Cytometry and MRC-LMS microscopy facility for support.

## Supplementary Figure legends

**Figure S1.**
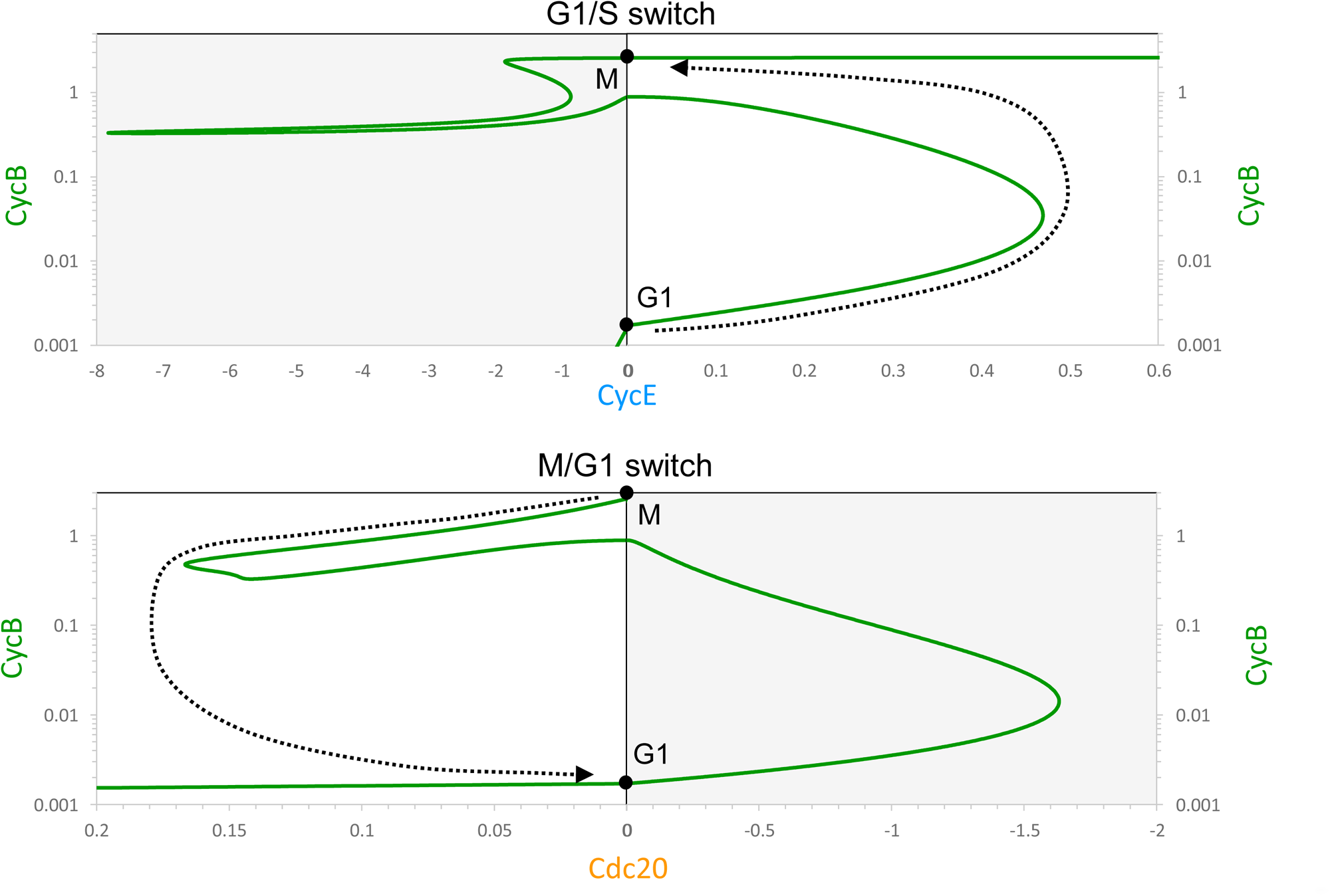
Bifurcation diagrams for CycB level as a function of CycE or Cdc20. See details in Fig. 2.

**Figure S2.**
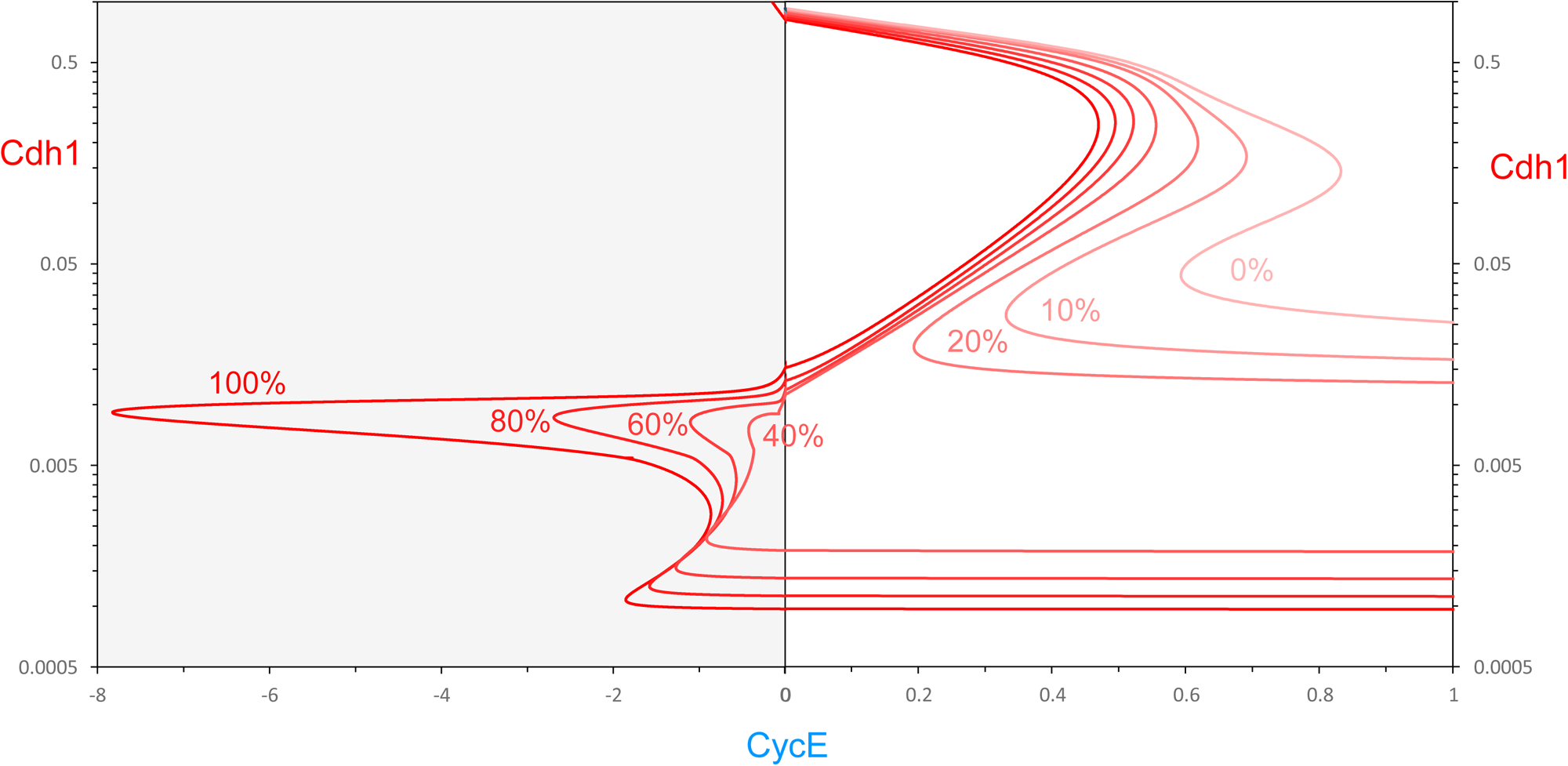
Bifurcation diagrams that account for Cdh1 endocycles. Cdh1 vs CycE bifurcation curves for increasing inhibition of Cdk1 activity. Percentage refers to level of Cdk1 activity remaining, i.e., 100% means full activity, 0% is no activity. If Cdk1 activity is < 25%, the gate at **M** (CycE = 0, Cdh1 low) fails to latch. As CycE level drops, Cdh1 will spontaneously reactivate.

**Figure S3.**
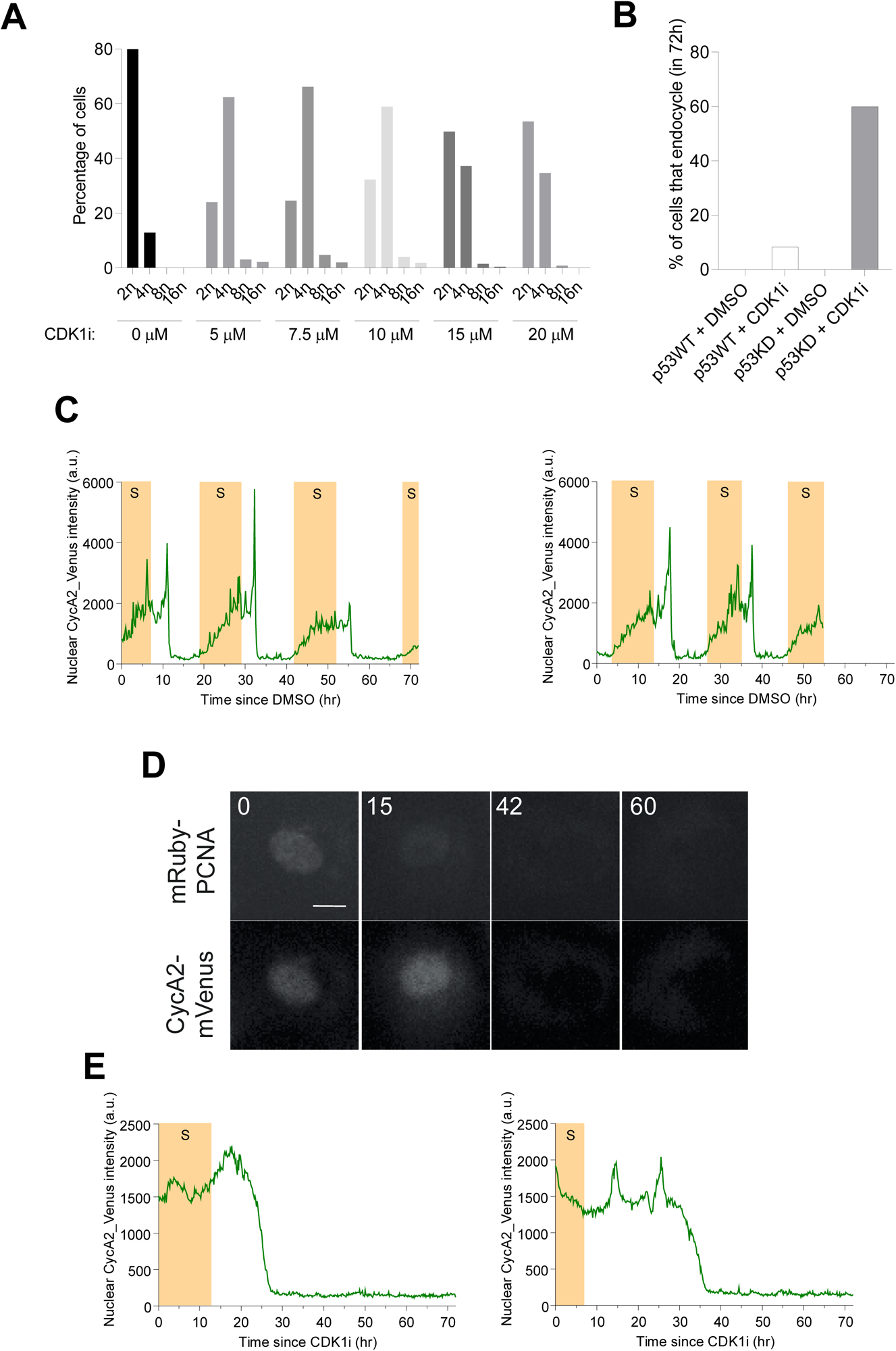
Cdk1 inhibition-induced endoreplication (Cdh1 endocycles). **(A)** The percentage of cells of different ploidy, as quantified by flow cytometry, after treatment with different doses of the Cdk1i, RO-3306, for 72 h. Discrete 8n and 16n peaks are characteristic of endoreplication. **(B)** The percentage of cells undergoing at least one endocycle in the 72 h imaging window in each condition. WT is wild-type p53. KD is p53 knockdown by siRNA. **(C)** Fluctuations of CycA2-mVenus during mitotic cycles in the presence of vehicle (DMSO). **(D)** CycA2-mVenus in individual cells that do not undergo endocycles. Still images of mRuby-PCNA and CycA2-mVenus labelled nuclei from timelapse experiments. Time shown in hours. Scale bar is 10 µm**. (E)** Graphs showing quantification of CycA2-mVenus, plotted from the time of CDK1i addition (t = 0 h). Shaded yellow areas represent S phase, as defined by mRuby-PCNA foci.

**Figure S4.**
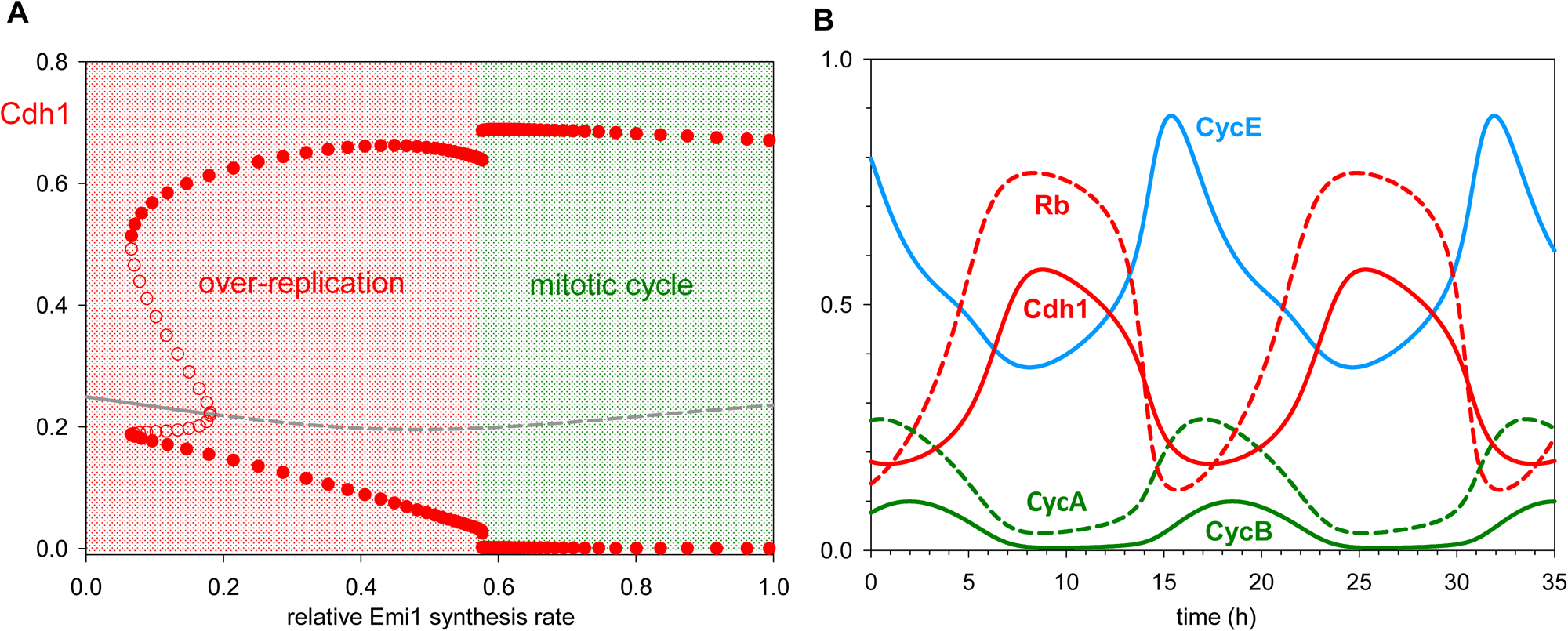
Down-regulation of Emi1 synthesis converts mitotic cycles into over-replication cycles. **(A)** Bifurcation diagram: Cdh1 activity as a function of the relative synthesis rate of Emi1. Solid (dashed) gray line: stable (unstable) steady states; solid (open) red circles: maximum and minimum excursions of Cdh1 activity during stable (unstable) limit cycle oscillations. Notice that Cdh1 axis is linear compared to logarithmic on Fig. 4A. (**B**) Simulation of Cdh1 endocycles for 90% suppression of Emi1 synthesis. Cdh1 activity never drops very low, so (presumably) replication origins are continuously relicensed, and DNA synthesis proceeds continuously rather than in discrete rounds of replication.

**Figure S5.**
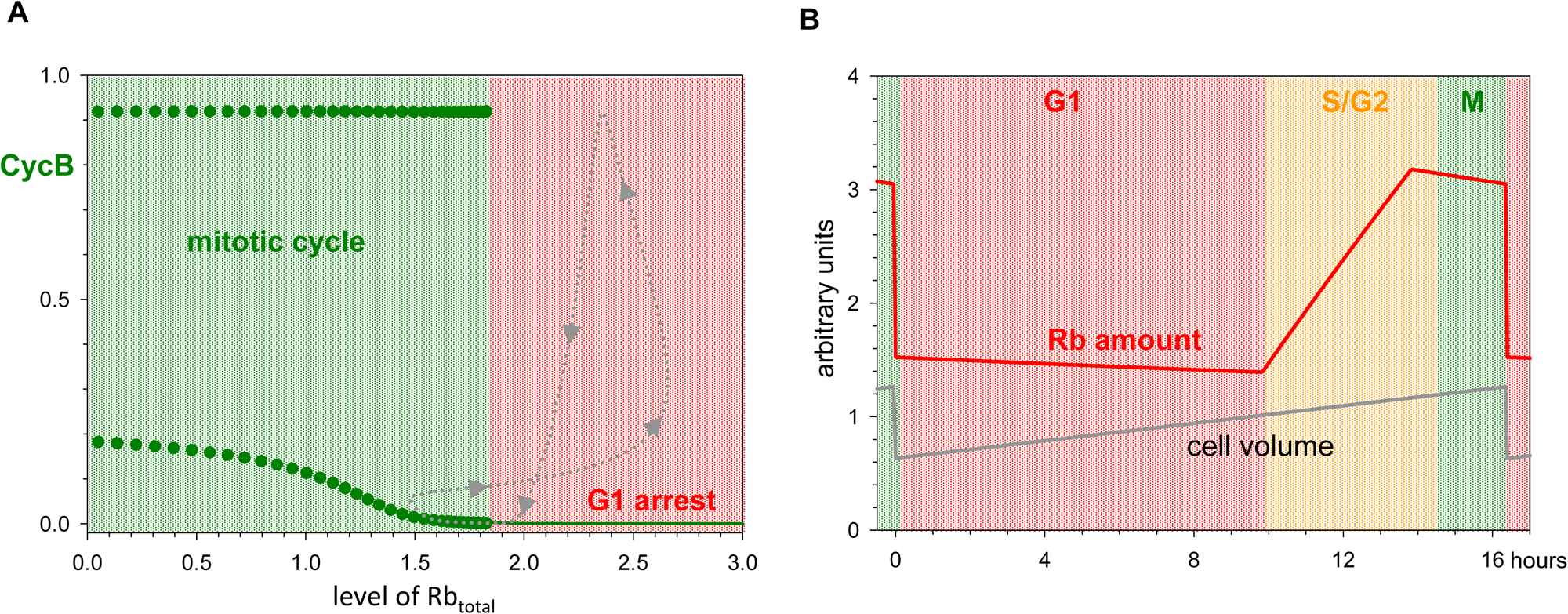
Checkpoint mechanisms convert spontaneous cell-cycle oscillations into conditional cycles, contingent on execution of certain events. **(A)** Bifurcation diagram (CycB vs concentration of Rb_total_) for the cell-growth checkpoint (G1/S transition). Solid green lines: stable steady states; solid green circles: maximum and minimum activity of CycB on a limit-cycle oscillation (spontaneous mitotic cycles). The dashed gray line is the conditional cell cycle, contingent on [Rb_total_] dilution by cell growth (movement from right to left) and the doubling of total Rb amount (movement from left to right) when the *retinoblastoma* protein is synthesized in S phase. The limit cycles arise at a ‘homoclinic saddle-loop’ bifurcation at [Rb_total_] ≈ 1.83. **(B)** Temporal changes of Rb amount and cell volume during the cell cycle in the simulation of Figure 6.

**Figure S6.**
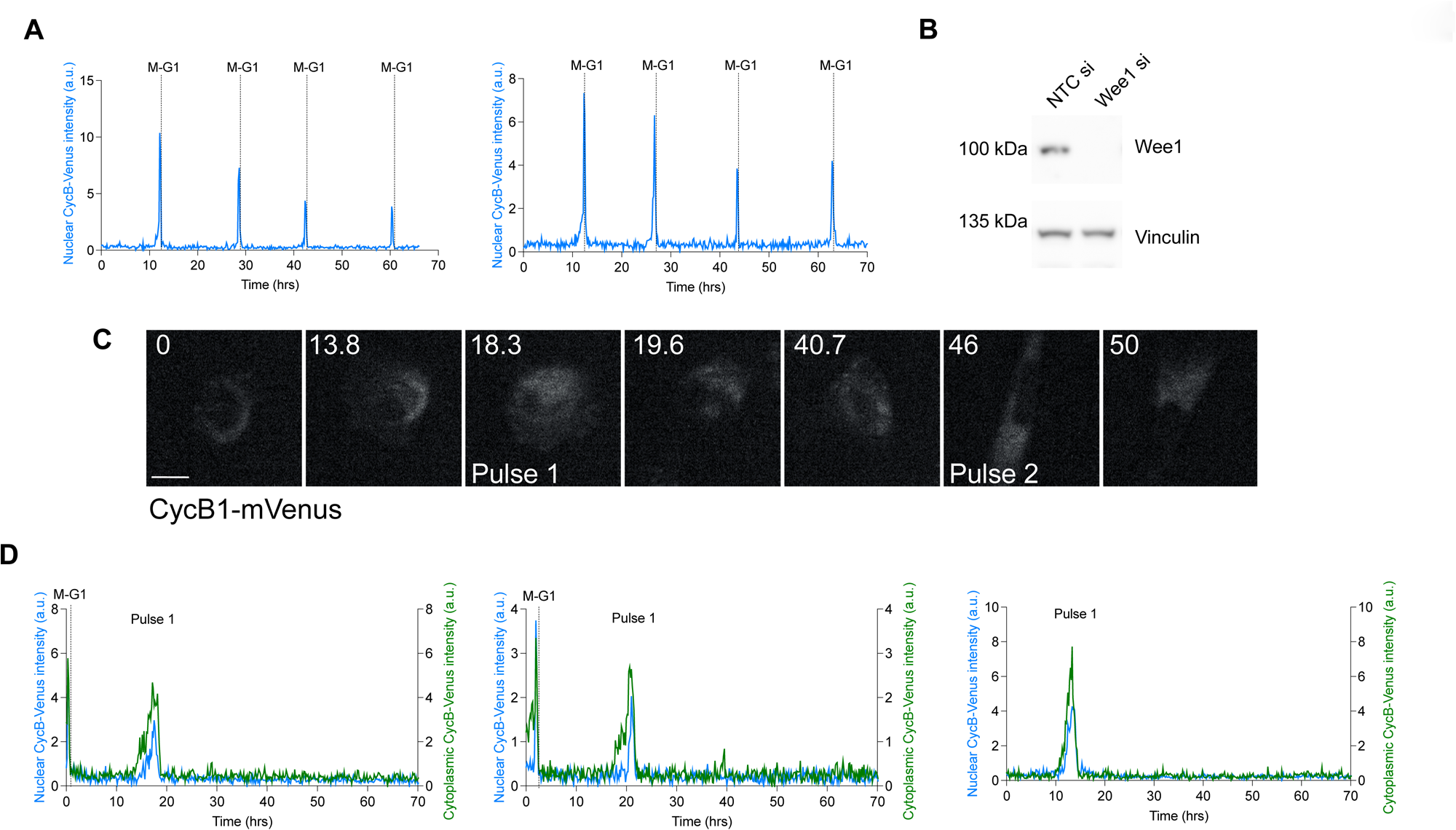
Constitutively active Cdk1:CycB induces Cdc20-endocycles. (**A)** Normalised CycB1-mVenus intensity in individual cells treated with control siRNA and undergoing normal mitotic cycles. **(B)** Western blot for Wee1 in NTC and Wee1 siRNA treated cells. Vinculin is used as a loading control. **(C)** Still images of hTert-RPE1 CycB1-mVenus labelled cells from timelapse experiments. Wee1 was depleted by siRNA 6 h prior to the start of filming (t = 0 h). The cell displays two interphase pulses of CycB1-mVenus expression in the absence of further mitoses. Time shown in hours. Scale bar is 10 µm. **(D)** Normalised CycB1-mVenus intensity in individual cells treated with Wee1 siRNA with one pulse only, plotted from the time of timelapse start (t = 0 h).

## References

1. Morgan, D.O. (2007). The Cell Cycle: Principles of Control. (New Science Press).

2. Pardee, A.B. (1974). A restriction point for control of normal animal cell proliferation. Proc Natl Acad Sci U S A 71, 1286–1290. 10.1073/pnas.71.4.1286.

3. Musacchio, A. (2015). The Molecular Biology of Spindle Assembly Checkpoint Signaling Dynamics. Curr Biol 25, R1002–1018. 10.1016/j.cub.2015.08.051.

4. Kohn, K.W. (1999). Molecular interaction map of the mammalian cell cycle control and DNA repair systems. Mol Biol Cell 10, 2703–2734. 10.1091/mbc.10.8.2703.

5. Chen, K.C., Csikasz-Nagy, A., Gyorffy, B., Val, J., Novak, B., and Tyson, J.J. (2000). Kinetic analysis of a molecular model of the budding yeast cell cycle. Mol Biol Cell 11, 369–391. 10.1091/mbc.11.1.369.

6. Tyson, J.J., and Novak, B. (2008). Temporal organization of the cell cycle. Curr Biol 18, R759–R768. 10.1016/j.cub.2008.07.001.

7. Novak, B., and Tyson, J.J. (2022). Mitotic kinase oscillation governs the latching of cell cycle switches. Curr Biol 32, 2780–2785 e2782. 10.1016/j.cub.2022.04.016.

8. Nasmyth, K. (1996). At the heart of the budding yeast cell cycle. Trends Genet 12, 405–412. 10.1016/0168-9525(96)10041-x.

9. Nurse, P. (1990). Universal control mechanism regulating onset of M-phase. Nature 344, 503–508. 10.1038/344503a0.

10. Clurman, B.E., Sheaff, R.J., Thress, K., Groudine, M., and Roberts, J.M. (1996). Turnover of cyclin E by the ubiquitin-proteasome pathway is regulated by cdk2 binding and cyclin phosphorylation. Genes Dev 10, 1979–1990. 10.1101/gad.10.16.1979.

11. Peters, J.M. (2006). The anaphase promoting complex/cyclosome: a machine designed to destroy. Nat Rev Mol Cell Biol 7, 644–656. 10.1038/nrm1988.

12. Hagting, A., Den Elzen, N., Vodermaier, H.C., Waizenegger, I.C., Peters, J.M., and Pines, J. (2002). Human securin proteolysis is controlled by the spindle checkpoint and reveals when the APC/C switches from activation by Cdc20 to Cdh1. J Cell Biol 157, 1125–1137. 10.1083/jcb.200111001.

13. Cappell, S.D., Mark, K.G., Garbett, D., Pack, L.R., Rape, M., and Meyer, T. (2018). EMI1 switches from being a substrate to an inhibitor of APC/C(CDH1) to start the cell cycle. Nature 558, 313–317. 10.1038/s41586-018-0199-7.

14. Fischer, M., Schade, A.E., Branigan, T.B., Muller, G.A., and DeCaprio, J.A. (2022). Coordinating gene expression during the cell cycle. Trends Biochem Sci 47, 1009–1022. 10.1016/j.tibs.2022.06.007.

15. Bertoli, C., Skotheim, J.M., and de Bruin, R.A. (2013). Control of cell cycle transcription during G1 and S phases. Nat Rev Mol Cell Biol 14, 518–528. 10.1038/nrm3629.

16. Vigneron, S., Sundermann, L., Labbe, J.C., Pintard, L., Radulescu, O., Castro, A., and Lorca, T. (2018). Cyclin A-cdk1-Dependent Phosphorylation of Bora Is the Triggering Factor Promoting Mitotic Entry. Dev Cell 45, 637–650 e637. 10.1016/j.devcel.2018.05.005.

17. Hansen, D.V., Loktev, A.V., Ban, K.H., and Jackson, P.K. (2004). Plk1 regulates activation of the anaphase promoting complex by phosphorylating and triggering SCFbetaTrCP-dependent destruction of the APC Inhibitor Emi1. Mol Biol Cell 15, 5623–5634. 10.1091/mbc.e04-07-0598.

18. Novak, B., and Tyson, J.J. (1993). Numerical analysis of a comprehensive model of M-phase control in Xenopus oocyte extracts and intact embryos. J Cell Sci 106 ( *Pt 4*), 1153–1168. 10.1242/jcs.106.4.1153.

19. Pomerening, J.R., Sontag, E.D., and Ferrell, J.E., Jr. (2003). Building a cell cycle oscillator: hysteresis and bistability in the activation of Cdc2. Nat Cell Biol 5, 346–351. 10.1038/ncb954.

20. Sha, W., Moore, J., Chen, K., Lassaletta, A.D., Yi, C.S., Tyson, J.J., and Sible, J.C. (2003). Hysteresis drives cell-cycle transitions in Xenopus laevis egg extracts. Proc Natl Acad Sci U S A 100, 975–980. 10.1073/pnas.0235349100.

21. Mochida, S., Maslen, S.L., Skehel, M., and Hunt, T. (2010). Greatwall phosphorylates an inhibitor of protein phosphatase 2A that is essential for mitosis. Science 330, 1670–1673. 10.1126/science.1195689.

22. Gharbi-Ayachi, A., Labbe, J.C., Burgess, A., Vigneron, S., Strub, J.M., Brioudes, E., Van-Dorsselaer, A., Castro, A., and Lorca, T. (2010). The substrate of Greatwall kinase, Arpp19, controls mitosis by inhibiting protein phosphatase 2A. Science 330, 1673–1677. 10.1126/science.1197048.

23. Cappell, S.D., Chung, M., Jaimovich, A., Spencer, S.L., and Meyer, T. (2016). Irreversible APC(Cdh1) Inactivation Underlies the Point of No Return for Cell-Cycle Entry. Cell 166, 167–180. 10.1016/j.cell.2016.05.077.

24. Ermentrout, B. (2002). Simulating, Analyzing, and Animating Dynamical Systems: A Guide to XPPAUT for Researchers and Students (Society for Industrial and Applied Mathematics).

25. Hayles, J., Fisher, D., Woollard, A., and Nurse, P. (1994). Temporal order of S phase and mitosis in fission yeast is determined by the state of the p34cdc2-mitotic B cyclin complex. Cell 78, 813–822. 10.1016/s0092-8674(94)90542-8.

26. Haase, S.B., Winey, M., and Reed, S.I. (2001). Multi-step control of spindle pole body duplication by cyclin-dependent kinase. Nat Cell Biol 3, 38–42. 10.1038/35050543.

27. Edgar, B.A., and Orr-Weaver, T.L. (2001). Endoreplication cell cycles: more for less. Cell 105, 297–306. 10.1016/s0092-8674(01)00334-8.

28. Itzhaki, J.E., Gilbert, C.S., and Porter, A.C. (1997). Construction by gene targeting in human cells of a ‘conditional’ CDC2 mutant that rereplicates its DNA. Nat Genet 15, 258–265. 10.1038/ng0397-258.

29. Ma, H.T., Tsang, Y.H., Marxer, M., and Poon, R.Y. (2009). Cyclin A2-cyclin-dependent kinase 2 cooperates with the PLK1-SCFbeta-TrCP1-EMI1-anaphase-promoting complex/cyclosome axis to promote genome reduplication in the absence of mitosis. Mol Cell Biol 29, 6500–6514. 10.1128/MCB.00669-09.

30. Laronne, A., Rotkopf, S., Hellman, A., Gruenbaum, Y., Porter, A.C., and Brandeis, M. (2003). Synchronization of interphase events depends neither on mitosis nor on cdk1. Mol Biol Cell 14, 3730–3740. 10.1091/mbc.e02-12-0850.

31. Zerjatke, T., Gak, I.A., Kirova, D., Fuhrmann, M., Daniel, K., Gonciarz, M., Muller, D., Glauche, I., and Mansfeld, J. (2017). Quantitative Cell Cycle Analysis Based on an Endogenous All-in-One Reporter for Cell Tracking and Classification. Cell Rep 19, 1953–1966. 10.1016/j.celrep.2017.05.022.

32. Mansfeld, J., Collin, P., Collins, M.O., Choudhary, J.S., and Pines, J. (2011). APC15 drives the turnover of MCC-CDC20 to make the spindle assembly checkpoint responsive to kinetochore attachment. Nat Cell Biol 13, 1234–1243. 10.1038/ncb2347.

33. Machida, Y.J., and Dutta, A. (2007). The APC/C inhibitor, Emi1, is essential for prevention of rereplication. Genes Dev 21, 184–194. 10.1101/gad.1495007.

34. Barr, A.R., Heldt, F.S., Zhang, T., Bakal, C., and Novak, B. (2016). A Dynamical Framework for the All-or-None G1/S Transition. Cell Syst 2, 27–37. 10.1016/j.cels.2016.01.001.

35. Pomerening, J.R., Ubersax, J.A., and Ferrell, J.E., Jr. (2008). Rapid cycling and precocious termination of G1 phase in cells expressing CDK1AF. Mol Biol Cell 19, 3426–3441. 10.1091/mbc.e08-02-0172.

36. Zatulovskiy, E., Zhang, S., Berenson, D.F., Topacio, B.R., and Skotheim, J.M. (2020). Cell growth dilutes the cell cycle inhibitor Rb to trigger cell division. Science 369, 466–471. 10.1126/science.aaz6213.

37. Collin, P., Nashchekina, O., Walker, R., and Pines, J. (2013). The spindle assembly checkpoint works like a rheostat rather than a toggle switch. Nat Cell Biol 15, 1378–1385. 10.1038/ncb2855.

38. Cross, F.R., Archambault, V., Miller, M., and Klovstad, M. (2002). Testing a mathematical model of the yeast cell cycle. Mol Biol Cell 13, 52–70. 10.1091/mbc.01-05-0265.

39. Lopez-Aviles, S., Kapuy, O., Novak, B., and Uhlmann, F. (2009). Irreversibility of mitotic exit is the consequence of systems-level feedback. Nature 459, 592–595. 10.1038/nature07984.

40. Simmons Kovacs, L.A., Mayhew, M.B., Orlando, D.A., Jin, Y., Li, Q., Huang, C., Reed, S.I., Mukherjee, S., and Haase, S.B. (2012). Cyclin-dependent kinases are regulators and effectors of oscillations driven by a transcription factor network. Mol Cell 45, 669–679. 10.1016/j.molcel.2011.12.033.

41. Lu, Y., and Cross, F.R. (2010). Periodic cyclin-Cdk activity entrains an autonomous Cdc14 release oscillator. Cell 141, 268–279. 10.1016/j.cell.2010.03.021.

42. Manzoni, R., Montani, F., Visintin, C., Caudron, F., Ciliberto, A., and Visintin, R. (2010). Oscillations in Cdc14 release and sequestration reveal a circuit underlying mitotic exit. J Cell Biol 190, 209–222. 10.1083/jcb.201002026.

43. Bielski, C.M., Zehir, A., Penson, A.V., Donoghue, M.T.A., Chatila, W., Armenia, J., Chang, M.T., Schram, A.M., Jonsson, P., Bandlamudi, C., et al. (2018). Genome doubling shapes the evolution and prognosis of advanced cancers. Nat Genet 50, 1189–1195. 10.1038/s41588-018-0165-1.

44. Dewhurst, S.M., McGranahan, N., Burrell, R.A., Rowan, A.J., Gronroos, E., Endesfelder, D., Joshi, T., Mouradov, D., Gibbs, P., Ward, R.L., et al. (2014). Tolerance of whole-genome doubling propagates chromosomal instability and accelerates cancer genome evolution. Cancer Discov 4, 175–185. 10.1158/2159-8290.CD-13-0285.

45. Lopez, S., Lim, E.L., Horswell, S., Haase, K., Huebner, A., Dietzen, M., Mourikis, T.P., Watkins, T.B.K., Rowan, A., Dewhurst, S.M., et al. (2020). Interplay between whole-genome doubling and the accumulation of deleterious alterations in cancer evolution. Nat Genet 52, 283–293. 10.1038/s41588-020-0584-7.

46. Quinton, R.J., DiDomizio, A., Vittoria, M.A., Kotynkova, K., Ticas, C.J., Patel, S., Koga, Y., Vakhshoorzadeh, J., Hermance, N., Kuroda, T.S., et al. (2021). Whole-genome doubling confers unique genetic vulnerabilities on tumour cells. Nature 590, 492–497. 10.1038/s41586-020-03133-3.

47. Fujiwara, T., Bandi, M., Nitta, M., Ivanova, E.V., Bronson, R.T., and Pellman, D. (2005). Cytokinesis failure generating tetraploids promotes tumorigenesis in p53-null cells. Nature 437, 1043–1047. 10.1038/nature04217.

48. Wolf, F., Wandke, C., Isenberg, N., and Geley, S. (2006). Dose-dependent effects of stable cyclin B1 on progression through mitosis in human cells. EMBO J 25, 2802–2813. 10.1038/sj.emboj.7601163.

49. Vazquez-Novelle, M.D., Sansregret, L., Dick, A.E., Smith, C.A., McAinsh, A.D., Gerlich, D.W., and Petronczki, M. (2014). Cdk1 inactivation terminates mitotic checkpoint surveillance and stabilizes kinetochore attachments in anaphase. Curr Biol 24, 638–645. 10.1016/j.cub.2014.01.034.

50. Bardin, A.J., and Amon, A. (2001). Men and sin: what’s the difference? Nat Rev Mol Cell Biol 2, 815–826. 10.1038/35099020.

51. Otto, T., and Sicinski, P. (2017). Cell cycle proteins as promising targets in cancer therapy. Nat Rev Cancer 17, 93–115. 10.1038/nrc.2016.138.

52. Hirai, H., Iwasawa, Y., Okada, M., Arai, T., Nishibata, T., Kobayashi, M., Kimura, T., Kaneko, N., Ohtani, J., Yamanaka, K., et al. (2009). Small-molecule inhibition of Wee1 kinase by MK-1775 selectively sensitizes p53-deficient tumor cells to DNA-damaging agents. Mol Cancer Ther 8, 2992–3000. 10.1158/1535-7163.MCT-09-0463.

53. Guiley, K.Z., Stevenson, J.W., Lou, K., Barkovich, K.J., Kumarasamy, V., Wijeratne, T.U., Bunch, K.L., Tripathi, S., Knudsen, E.S., Witkiewicz, A.K., et al. (2019). p27 allosterically activates cyclin-dependent kinase 4 and antagonizes palbociclib inhibition. Science 366. 10.1126/science.aaw2106.

